# Striatal dopamine reflects individual long-term learning trajectories

**DOI:** 10.1101/2023.12.14.571653

**Authors:** Samuel Liebana Garcia, Aeron Laffere, Chiara Toschi, Louisa Schilling, Jacek Podlaski, Matthias Fritsche, Peter Zatka-Haas, Yulong Li, Rafal Bogacz, Andrew Saxe, Armin Lak

## Abstract

Learning from naïve to expert occurs over long periods of time, accompanied by changes in the brain’s neuronal signals. The principles governing behavioural and neuronal dynamics during long-term learning remain unknown. We developed a psychophysical visual decision task for mice that allowed for studying learning trajectories from naïve to expert. Mice adopted sequences of strategies that became more stimulus-dependent over time, showing substantial diversity in the strategies they transitioned through and settled on. Remarkably, these transitions were systematic; the initial strategy of naïve mice predicted their strategy several weeks later. Longitudinal imaging of dopamine release in dorsal striatum demonstrated that dopamine signals evolved over learning, reflecting stimulus-choice associations linked to each individual’s strategy. A deep neural network model trained on the task with reinforcement learning captured behavioural and dopamine trajectories. The model’s learning dynamics accounted for the mice’s diverse and systematic learning trajectories through a hierarchy of saddle points. The model used prediction errors mirroring recorded dopamine signals to update its parameters, offering a concrete account of striatal dopamine’s role in long-term learning. Our results demonstrate that long-term learning is governed by diverse yet systematic transitions through behavioural strategies, and that dopamine signals exhibit key characteristics to support this learning.

## Main

Over long periods of time, individuals learn to make increasingly accurate choices in response to stimuli they perceive. For instance, after extended training, a naïve individual can learn to play tennis expertly. Past studies have argued that humans and animals often pass through several stages of behaviour as they learn (1). This long-term learning rarely occurs in lockstep. Rather, individuals often take various paths through the space of behaviour, resulting in diverse learning trajectories. Neuroscientific studies of learning have typically ignored the dynamics of these diverse learning trajectories. They have either investigated neuronal signals in expert animals as they fine-tune an already learned task, or contrasted neuronal signals between naïve and expert animals without consideration of their learning trajectories (2–13). As such, the behavioural, neural, and computational underpinnings of long-term learning trajectories remain unknown.

Long-term learning often occurs by trial and error, a process well-captured by reinforcement learning (RL) models (14). Dopaminergic neurons of the midbrain play an essential role in reinforcement learning, signalling reward prediction error (RPE), the difference between predicted and acquired reward (15,16). This RPE can be used to improve reward predictions, and therefore, future choices. However, little is known about the relationship between dopamine (DA) signals and the dynamics of long-term learning. Studies relating DA signals and learning have either measured these signals during Pavlovian conditioning, or during trial-by-trial decision making after task acquisition (17–30). It therefore remains unclear whether dopamine-driven learning can account for individual long-term learning trajectories.

Here we characterise individual behaviour as mice learned a perceptual decision making task over several weeks, while longitudinally measuring DA release in dorsolateral striatum (DLS), a region of the basal ganglia implicated in reinforcing reward-associated choices (31,32). Mice improved their performance by adopting sequences of strategies that became more stimulus-dependent over time. This sequence varied widely across individuals. However, the strategy transitions were remarkably systematic; each animal’s future behaviour could be predicted days in advance based on its current behaviour. DLS DA signals reflected each individual’s evolving behavioural strategies, showing strikingly similar patterns of diverse yet systematic transitions. DA responses to stimuli developed reflecting stimulus-choice associations. Importantly, this encoding was dissociated from choice accuracy; DA responses were absent for stimuli that predicted reward but did not inform choice. DA outcome signals mirrored the stimulus responses, decreasing in magnitude as DA responses to stimuli grew. A simple computational model based on reinforcement learning in a deep neural network effectively reproduced the observed distribution of behavioural and DA trajectories throughout learning. This model had two important features: depth, i.e. three layers (necessary for explaining learning trajectories), and teaching signals defined based on subsets of inputs (necessary to account for DA signals). An analytical description of the model’s learning dynamics revealed a hierarchy of saddle points that accounted for the systematic transitions of behaviour and DA signals from naïve to expert. These results suggest a model in which long-term learning traverses a continuous space of behaviours by acquiring stimulus-choice associations while greedily improving reward prediction, driven by dopamine-based learning signals.

### Long-term learning trajectories from naïve to expert

To study long-term learning of perceptual decision making, we further developed an established visual decision task in head-fixed mice (33). In each trial, we presented a visual stimulus (a grating) on the left or right side of a screen. The mouse, head-fixed in front of the screen, reported the stimulus position (left or right) by steering a wheel with its forepaws to bring the stimulus to the centre of the screen (Fig.1a, Extended Data Fig.1a, Methods). The mouse received a drop of water reward for each correct choice, or a white noise sound and brief timeout for incorrect choices (Fig.1a). The contrast of the grating stimulus changed across trials, making the trials easier or harder. Some trials did not have a stimulus (i.e. zero-contrast trials), for which animals were rewarded randomly regardless of the wheel movement direction (Fig.1a). We trained the mice over multiple days with a single session (approximately 250 trials) each day. Importantly, we kept the task unchanged throughout the entire experiment, presenting the full set of stimuli, task contingencies and trial timing from day one until expert performance (Fig.1a, Extended Data Fig.1a, Methods). This ensured that any changes in behaviour are a consequence of animals’ internal learning mechanisms, rather than a consequence of experimentally imposed changes to the task. It also enabled us to directly compare behavioural and neural data across days. Thus, the training differed from conventional shaping curricula in which the task becomes incrementally harder to guide learning. Mice learned the task reaching accuracies of at least 70% over many days of training (Fig.1b, Extended Data Fig.1b,c, median days = 19, median trial number = 3430), showing progressively steeper psychometric curves (Fig.1c), and faster choice response times (Extended Data Fig.1d). The decrease in response times (RT) occurred before the increase in the choice accuracy (Extended Data Fig.1e,f), leading to an early increase in reward rate (i.e., reward per unit time) while the accuracy was still at chance level (Extended Data Fig.1g).

**Fig 1.**
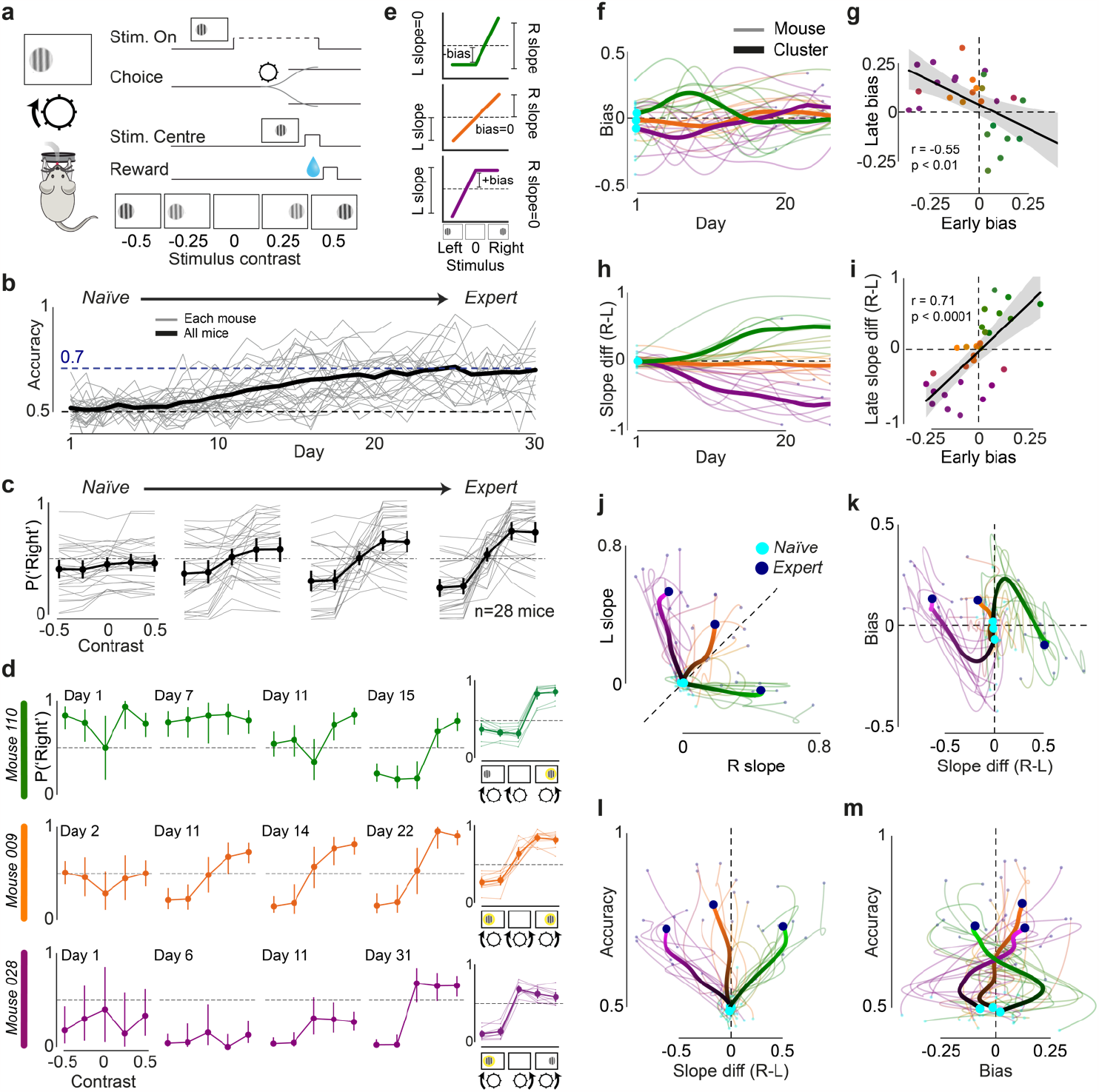
Long-term learning trajectories are diverse yet systematic. **a**, The visual decision task for head-fixed mice (*n*=28 mice; Methods). **b**, Accuracy over days per mouse (grey) and averaged across all mice (black). Dashed lines indicate chance level (black) and 70% accuracy level (blue). Number of days limited to 30 for better visualisation. **c**, Psychometric curve over quartiles per mouse (grey) and averaged across all mice (black). Quartiles are defined per mouse by dividing days into 4 groups, with any remainder added to the last group. Negative (positive) contrast values indicate stimuli presented on the left (right) side of the screen. P(‘Right’) indicates the probability of reporting ‘right’ side stimulus position (i.e., turning the wheel anticlockwise). Error bars indicate the 95% confidence interval across mice. **d**, Left, psychometric curves from 3 example mice on 4 example days throughout learning. Error bars indicate the 95% confidence interval of a two-sided binomial test on P(‘Right’). See Extended Data Fig.2a for corresponding chronometric curves. Right, per mouse (thin) and average expert psychometric curves clustered by trajectory type (thick): right-associating (green), balanced (orange) and left-associating (purple). See Extended Data Fig.2b for corresponding chronometric curves. Cluster labels for each mouse were obtained from panel j, and colours were obtained from Extended Data Fig.2c. Error bars indicate 95% confidence interval across mice in each cluster. **e**, Schematic explaining the metrics used to study behaviour. Left (right) slope is defined as the absolute difference between the P(‘Right’) for left (right) stimulus and zero-contrast trials. Bias is defined as the difference between zero-contrast P(‘Right’) and balanced choice probability (i.e., 0.5), thus indicating the choice preference on zero-contrast trials. **f**, Bias over days per mouse (thin) and for the 3 clusters from panel j (thick). In right- and left-associating mice, biases increase before reversing (*p*<0.05; t-test first 2 days vs. days 5-6). Number of days limited to 25 for better visualisation. **g**, Regression of early bias (average across days 4-8) against late bias (average across final 5 days). Each point represents a mouse. *p*-value calculated from the exact distribution of r (stats.scipy.pearsonr). **h**, Difference between right and left psychometric slopes over days per mouse (thin) and for the 3 clusters from panel j (thick). Number of days limited to 25 for better visualisation. **i**, Regression of early bias (average across days 4-8) against late slope difference (average across final 5 days). Each point represents a mouse. *p*-value from the exact distribution of r (stats.scipy.pearsonr). **j**, Left vs. right slope across days per mouse (thin) and for 3 clusters (thick). Cyan dots indicate the beginning of learning and navy dots indicate the end. The hue of the cluster lines changes from dark to light indicating progress through learning. Left-associating (purple), balanced (orange) and right-associating (green) clusters are obtained from dynamic time warping clustering (Methods) applied to the mouse trajectories. The clusters from this analysis are used in all other panels. **k-m**, In order, difference in right and left (R-L) slope vs. bias; R-L slope vs. accuracy and bias vs. accuracy across days per mouse (thin) and for the 3 clusters from panel j (thick).

In early days of learning, mice favoured left or right choices to varying degrees, exhibiting flat, but often biased, psychometric curves (Fig.1d, first column, Fig.1f,g). The biases often increased in these early days (Fig.1f), while RTs decreased (Extended Data Fig.1d,e). During this initial period, psychometric slopes were often close to zero, i.e. flat psychometric curves (Fig.1c,d). Thus, initial days were marked by strategies where the mice ignored the position of the visual stimulus for making choices, showing increasing biases with different directions across animals.

During later days of learning, mice’s choices began to depend on the location of visual stimuli, resulting in psychometric curves with increasing slopes. Importantly, slopes often developed differently for left and right stimuli, with a vast diversity across mice (Fig.1h-l). To visualise this diversity, we coloured each mouse’s learning trajectory based on the asymmetry of its psychometric slopes (Extended Data Fig.2c, Methods). While the diversity of slopes formed a continuum across mice (Fig.1j), to better visualise the main trends we clustered slope trajectories over learning (Methods). In some mice the slopes increased similarly on both sides, resulting in more balanced psychometric curves (Fig.1j). This indicates that their strategy involved associating both left and right stimuli with their corresponding choice directions (Fig.1d middle row). However, in several other mice, the slope primarily increased on one side while the other side remained flat, forming a one-sided psychometric curve (Fig.1j). These mice therefore solved the task by associating stimuli on one side of the screen with their corresponding choice direction, and making the alternative choice in trials in which the associated stimulus was absent (Fig.1d top and bottom rows). Consistent with this strategy, choices in zero-contrast trials and trials with non-associated stimuli were indistinguishable, i.e., the psychometric curve was flat on one side (Fig.1d, right column, Extended Data Fig.2d). These slope asymmetries often persisted as learning progressed (Fig.1l, Extended Data Fig.2f,g), and their corresponding signatures were observed in RTs as well as eye movements (Extended Data Fig.2b,e,h). Importantly, despite strong asymmetries in slopes, one-sided left- and right-associating animals reached similar levels of choice accuracy compared to balanced animals (Fig.1l). This is because both the presence and absence of stimuli can inform correct choices in the task, as there is only one stimulus per trial. Overall, we observed that in later days mice transitioned to more stimulus-dependent strategies, while exhibiting diversity in their use of left and right stimuli to make choices.

Behavioural transitions throughout learning were systematic. The bias in initial days strongly predicted the biases and psychometric curves in later stages of learning (Fig.1g,i). Mice with initial left bias developed a larger slope on the left side of their psychometric curve, whereas mice with initial right bias developed a larger slope on the right side (Extended Data Fig.2g). To achieve high accuracy despite asymmetric slopes, mice with more one-sided strategies reversed their initial bias during learning (Fig.1g,k,m). Thus, naïve mice started with varying levels of bias, and this initial bias determined which stimuli were associated with choices in later stages of learning.

Taken together, the results show that in learning to make visual decisions from naïve to expert, mice exhibited diverse learning trajectories involving systematic transitions through behavioural strategies.

### Dorsal striatal dopamine signals from naïve to expert

We recorded DA release from day one until expert behaviour in the dorsolateral striatum (DLS). We injected GRAB-DA (34) in the DLS of wild-type mice, implanted one or two optic fibres dorsal to the injection site (one for each hemisphere), and imaged DLS DA release every day during learning (Fig.2a,b, Extended Data Fig.3a; see Methods).

**Fig 2.**
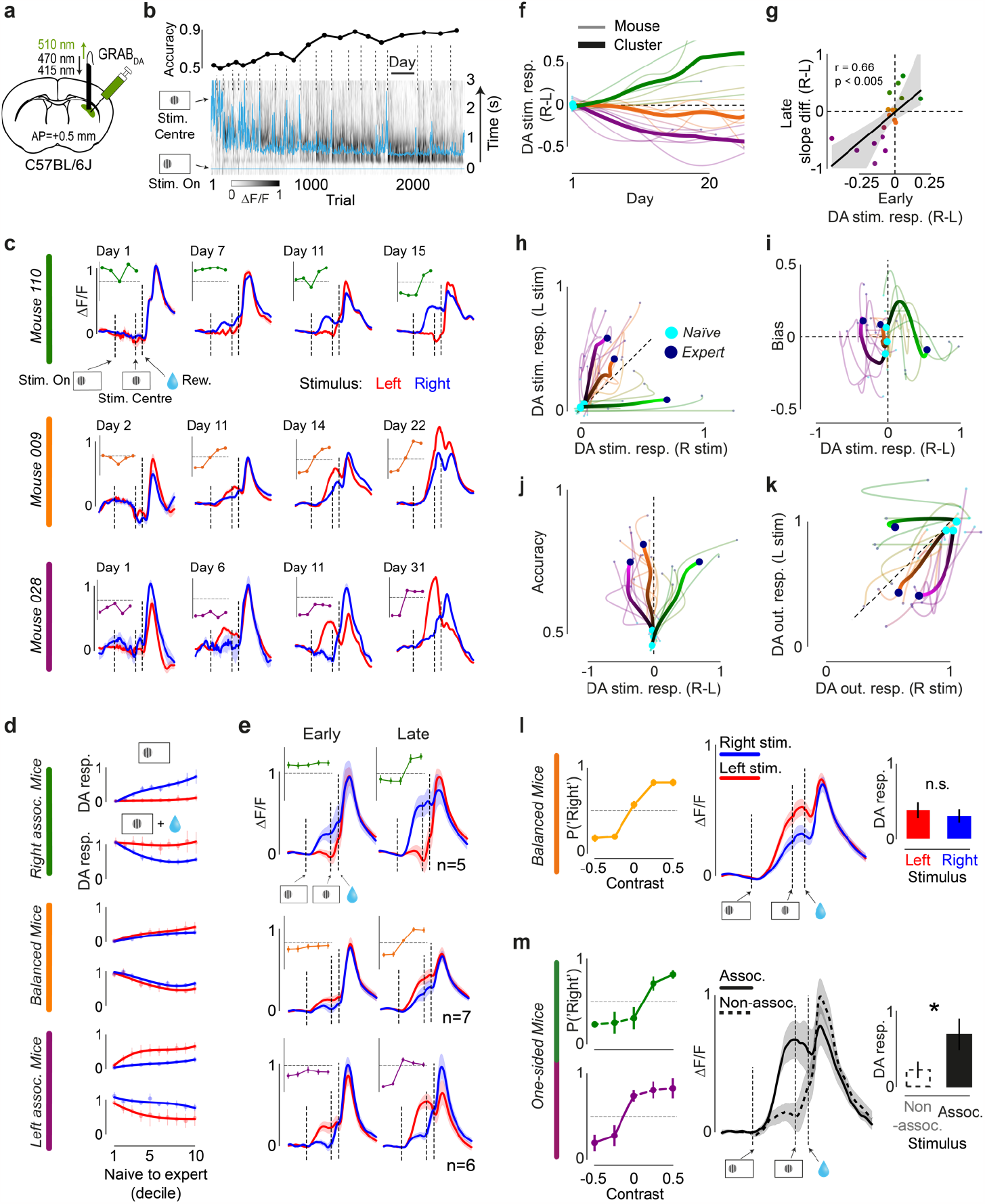
DLS DA encodes stimulus-choice association and reflects learning trajectories. **a**, Fibre photometry set-up for recording dopamine (DA) release in dorsolateral striatum (DLS). Recorded DA levels were normalised to correct for non-task-relevant day-by-day variations in fluorescence, see Methods. **b**, Accuracy over days and simultaneous trial-wise stim.-aligned DA recordings from an example mouse (only correct trials). Blue lines indicate 5-trial moving average of the time when the stimulus is brought to the centre (i.e., choice completion, top) and stimulus onset time (bottom). **c**, Average time warped DA signals in correct trials with stimulus on the left (red) and right (blue) for the same days and mice as in Fig.1d (psychometric curves reproduced as insets). Vertical dashed lines indicate stim. onset, stim. centre, and reward delivery time. Error bars indicate +/-s.e.m. across trials. **d**, Average stimulus and outcome DA responses over deciles in correct trials with stimulus on the left (red) and right (blue) for the three clusters from Fig.1j. Stim. and outcome responses were calculated by computing the maximum of smoothened fluorescence in an analysis window (see Methods). Outcome responses were defined as the sum of responses to stim. centre and reward delivery. Error bars indicate 95% confidence interval of average across mice. Data points fit with a 3rd degree spline to visualise trends (scipy.interpolate.UnivariateSpline). **e**, Left, average time warped DA signals in correct trials with stimulus on the left (red) and right (blue) and average psychometric curves for the three clusters from Fig.1j in early days (accuracy n.s. greater than 0.5). Right, average time warped DA signals and psychometric curves in late days (accuracy n.s. smaller than 0.7). Error bars indicate s.e.m. across mice. Day-by-day regression analyses confirmed these results (Extended Data Fig.6c). **f**, Difference in DA responses to right and left stimuli (R-L) over days per mouse (thin) and for the 3 clusters from Fig.1j (thick). Number of days limited to 25 for better visualisation. **g**, Regression of early difference in DA responses to R-L stimuli (average across days 4-8) against late R-L slope difference (average across final 5 days). Each point represents a mouse. *p*-value is calculated from the exact distribution of r. **h-k**, In order, right vs. left DA stim. responses; difference in DA responses to right and left stimuli (R-L) vs. bias; difference in DA responses to right and left stimuli (R-L) vs. accuracy; and DA rewarded outcome responses after right stimulus vs. after left stimulus per mouse (thin) and for the 3 clusters from Fig.1j. Cyan dots indicate the beginning of learning and navy dots indicate the end. The hue of the cluster lines changes from dark to light indicating progress through learning. **l**, Analysis of balanced mice DA responses in days where the difference in left and right stimulus trial choice accuracy is <0.1; Left, average psychometric curve from matched accuracy days. Middle, average time warped DA signals for correct trials with stimulus on the left (red) and right (blue). Right, bar plot showing the average DA responses to stimuli for correct left and right stim trials; *p* > 0.1, estimated using two-sided t-test. Error bars indicate 95% confidence interval of average across days. **m**, Same matched accuracy analysis as in Fig.2l but applied to left- and right-associating mice. * indicates *p <* 0.05.

**Fig 3.**
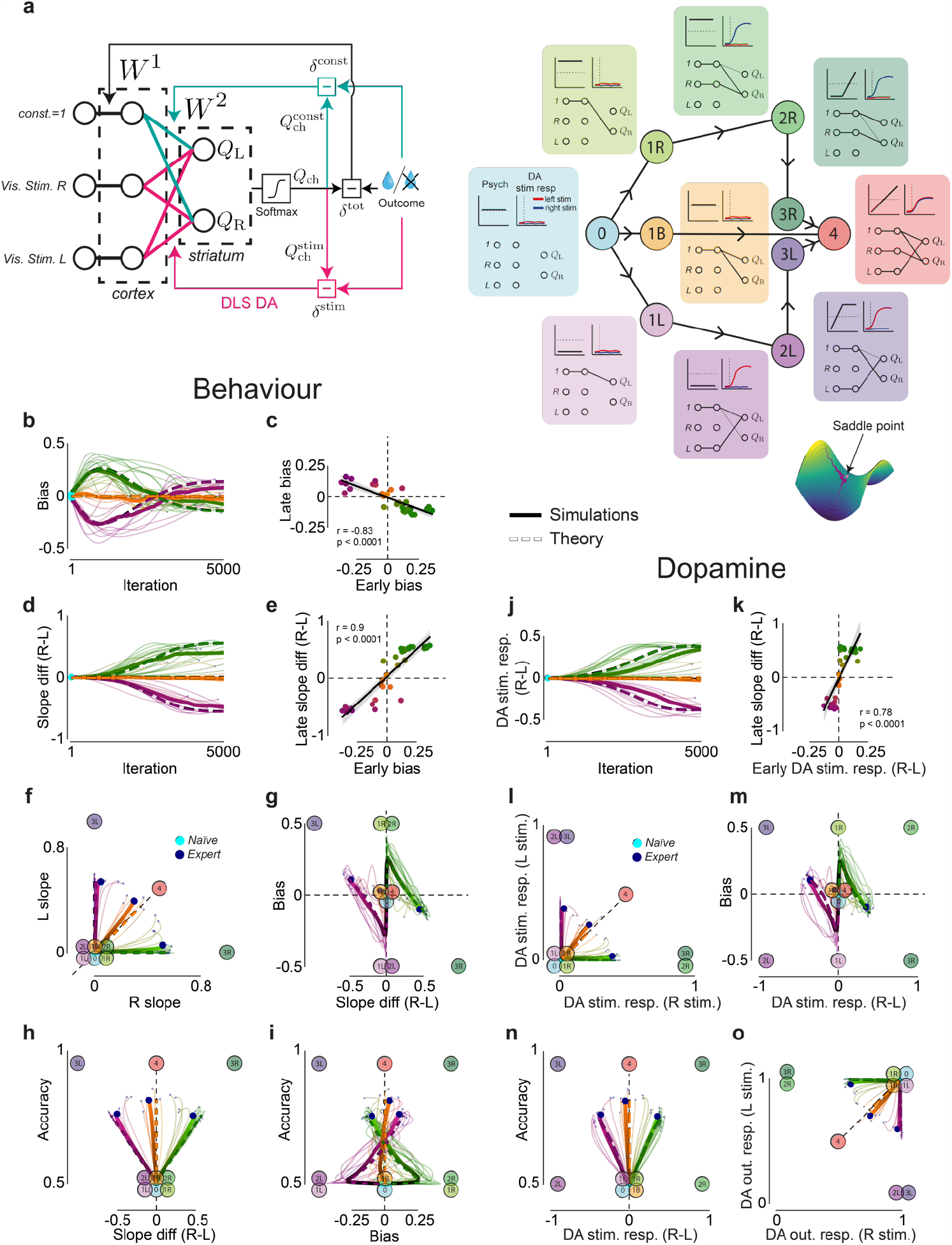
A deep linear neural network model accounts for behaviour and dopamine signals throughout learning. **a**, Left, schematic of the deep linear ‘tutor-executor’ RL network architecture and learning rule (Methods). Right, schematic of the stationary point structure including behavioural and neural predictions as well as corresponding network weight configurations. The connecting lines with arrows represent the steepest heteroclinic orbits into/out of each stationary point (Methods). All the stationary points are saddle points except for 4, which is the global minimum. **b**, c.f. Fig.1f, Bias over iterations per simulation (thin), for the 3 clusters from panel f (thick), and for the average dynamics (thick dashed). Here, and in panels d and j, the number of iterations is limited to 5000 for better visualisation. Thick dashed lines in all panels indicate trajectories derived from the analytical average dynamics (Methods). **c**, c.f. Fig.1g, Regression of early bias (average across iterations 1000-2000) against late bias (average across final 1000 iterations). Each point represents a simulation. *p*-value is calculated from the exact distribution of r. **d**, c.f. Fig.1h, Difference between right and left psychometric slopes over iterations per simulation (thin) and for the 3 clusters from panel f (thick). **e**, c.f. Fig.1i, Regression of early bias (average across iterations 1000-2000) against late slope difference (average across final 1000 iterations). Each point represents a simulation. *p*-value is calculated from the exact distribution of r. **f**, c.f. Fig.1j, Left vs. right slope over iterations per simulation (thin) and for 3 clusters (thick). Clusters and colours obtained using the same procedure as for the behavioural data in Fig.1j. The clusters from this analysis are used in all other panels. Stationary points here and in panels g-i are plotted using the average behaviour arising from their weight configurations. **g-i**, c.f. Fig.1k-m, in order, difference in right and left (R-L) slope vs. bias, R-L slope vs. accuracy, and bias vs. accuracy over iterations per simulation (thin) and for the 3 clusters from panel f (thick). **j**, c.f. Fig.2f, Difference in DA responses to right and left stimuli (R-L) over iterations per simulation (thin) and for the 3 clusters from panel f (thick). **k**, c.f. Fig 2g, Regression of early difference in DA responses to R-L stimuli (average across iterations 1000-2000) against late slope difference (average across final 1000 iterations). Each point represents a simulation. *p*-value is calculated from the exact distribution of r. **l-o**, c.f. Fig.2h-k, in order, right vs. left DA stim. responses; difference in DA responses to right and left stimuli (R-L) vs. bias; difference in DA responses to right and left stimuli (R-L) vs. accuracy; and DA rewarded outcome responses after right stimulus vs. after left stimulus per simulation (thin) and for the 3 clusters from panel f. Stationary points are plotted using the average DA responses arising from their weight configurations.

DA release occurred in response to specific events within a trial, and changed throughout learning. A kernel regression revealed that the onset of visual stimulus, the arrival of the stimulus to the centre of the screen in correct trials (i.e., ‘stim. centre’), and water reward (or its absence in error trials) were the main events modulating DA release (Extended Data Fig.3b,c). For our analyses, we defined a DA ‘stimulus’ and ‘outcome’ response to examine changes over learning. The ‘stimulus’ response quantified DA release levels in a short time window after stimulus onset (Methods). The ‘outcome’ response was defined as the sum of DA responses to completion of choice (marked by the stimulus arriving to the centre of the screen in the case of correct choices, or by the stimulus leaving the screen in the case of incorrect choices) and water reward delivery (or its absence). This is because the final stimulus position determines whether water reward will be subsequently delivered. In initial days, DA release mostly occurred in response to rewarded outcomes but not visual stimuli. As learning progressed, DA responses to visual stimuli grew and DA responses to water rewards diminished (Fig.2b-e, Extended Data Fig.3b,c, Extended Data Fig.4a-c). In error trials, when reward was not delivered, DA signals transiently decreased (Extended Data Fig.3b).

The development of DA responses during learning mirrored the diverse learning trajectories observed in behaviour. DA responses in individual mice showed strong correspondence to the development of their psychometric curves (Fig.2c, Extended Data Fig.4a). In mice developing more one-sided psychometric curves, DA responses to stimuli emerged most strongly for stimuli presented on the associated side (top and bottom rows in Fig.2c,d, Extended Data Fig.4a,b). However, in mice developing more balanced psychometric curves, DA responses to stimuli presented on left and right sides were similar (middle rows in Fig.2c,d and Extended Data Fig.4a,b). The diverse behavioural trajectories were also reflected in the rewarded outcome DA response; these DA responses were small after associated stimuli and large after non-associated stimuli (Fig.2c,d, Extended Data Fig.4a,b).

In early days, when psychometric curves were still flat, DA responses to stimuli were already appearing, developing differently for stimuli on the left and right side depending on the animal’s bias (Fig.2e Left). These DA responses were strongest for stimuli appearing on the same side as the choice bias (Fig.2e Left, Extended Data Fig.4c). We asked whether these signals reflected associations that the animals were forming between stimuli and choices, or whether they simply reflected each animal’s choice side preference. To address this, we inspected trials without visual stimuli (zero-contrast trials), and did not observe a difference in DA responses for the two choices (Extended Data Fig.4d). This indicated that these DA signals are a signature of the association animals were forming between stimuli and choices in early days, which also manifested in a decreased RT for choices after associated stimuli (Extended Data Fig.4e). Moreover, DA responses to stimuli were not evident on the first day of the experiment (Extended Data Fig.4f), indicating that they emerge over learning, rather than reflecting stimulus novelty (23,17,35). The early DA stimulus responses predicted both future DA responses to stimuli (Fig.2f, Extended Data Fig.4g,h) as well as the shape of psychometric curves in later days (Fig.2g). DA signatures of associations being formed between stimuli and choices were also evident in DA responses at trial outcome (Fig.2d, Extended Data Fig.4b,c,h,i). Thus, during initial days, DA responses emerged, reflecting the first signature of learning to associate stimuli and choices, and these DA signals predicted behaviour and DA signals many days later.

In later days, as the psychometric curves developed, DA responses strongly reflected the growing psychometric slopes observed across animals (Fig.2c-f). In right-associating mice (i.e. those with a steep psychometric curve on the right and a flat curve on the left), DA responses to stimuli were evident in response to right but not left stimuli (Fig.2e, top row). This pattern was opposite in left-associating mice (Fig.2e, bottom row). In balanced mice (i.e., those associating both stimuli with their corresponding choices), both left and right stimuli elicited strong DA responses (Fig.2e, middle row). These observations held independent of stimulus laterality with respect to the recorded brain hemisphere (Extended Data Fig.5a,b). The relationship between psychometric slopes and DA responses to stimuli was maintained even after controlling for any differences in the accuracy of left and right choices, i.e., selecting days where the accuracies were matched but the slopes differed (Fig.2l,m, Extended Data Fig.5c-e). Thus, in left- and right-associating mice, DA responses to associated stimuli were significantly larger than to non-associated stimuli, despite equally high choice accuracy for both stimuli (Fig.2m). This indicates that the DA responses correlate with the psychometric slope, i.e. the formed association between the stimulus and choice, rather than reflecting the choice accuracy. Moreover, the encoding of psychometric slope was largely invariant to RTs (Extended Data Fig.5f) and the accuracy of pending choice (i.e., correct/error, Extended Data Fig.5g). DA outcome responses reflected the difference between outcome value and the learned stimulus-choice association, i.e., the reward that animals learned to predict using each stimulus. Hence, these DA responses decreased as stimulus-choice associations further increased (Fig.2d, Extended Data Fig.5g, middle row), instead of reflecting choice accuracy (Fig.2m).

The dynamics of DA signals throughout the entirety of learning showed striking similarities to behavioural trajectories (Fig.2h-k, neural trajectories plotted with colours and clusters obtained from behavioural trajectories). DA responses to stimuli developed across days reflecting the dynamics of slope and bias that each animal exhibited during learning (Fig.2h-j). Similarly, DA responses to rewards developed over learning mirroring the evolving stimulus responses (Fig.2k, Extended Data Fig.6a,b). Thus, from DA signals at any point during learning, it was possible to infer the animals’ past and future DA signals and behavioural strategies (Fig.2f-k, Extended Data Fig.4g-i, and Extended Data Fig.6a,b).

Taken together, these results show that during long-term learning to make visual decisions, DLS DA responses to stimuli emerge as strategies become more stimulus-dependent, encoding stimulus-choice associations and predicting individuals’ learning trajectories. DLS DA outcome responses encode the difference between the value of trial outcome and reward predictions based on the visual stimulus, thus reflecting individuals’ learning trajectories.

### A deep linear neural network model of long-term learning

To understand the principles underlying the learning trajectories and their dopaminergic correlates, we designed a deep linear neural network model. In essence, this model contains multiple layers of neurons that learn to predict the reward associated with each stimulus and choice. The model has a simple architecture: an input layer of three neurons, a hidden layer of three neurons, and an output layer of two neurons (Fig.3a Left, Methods). A matrix of nonnegative weights denoted by *W*^*1*^ connects the input layer to the hidden layer, and similarly nonnegative weights in *W*^*2*^ connect the hidden layer to the output layer. The weights in *W*^*1*^ form independent channels from the input layer neurons to the ‘cortical’ neurons in the hidden layer, resembling the anatomical segregation of visual inputs in each brain hemisphere. The weights in *W*^*2*^ represent the brain’s ‘cortico-striatal’ synapses and determine the contribution of neurons in the hidden layer to the output. The outputs of the network are two action values, *Q*_*L*_ and *Q*_*R*_, reflecting the learned value (i.e., reward prediction) of taking each choice as a function of the inputs.

On each trial, the model receives inputs, makes a choice, and learns from the resulting outcome. The sensory inputs are binary values (represented as 0/1). Two inputs ‘*Vis. stim. R*’ and ‘*Vis. stim. L*’ indicate which of the stimuli is presented on a particular trial. The third input ‘*constant*’ is always set to 1 to reflect stimulus-independent input, capturing environmental features that do not change trial-by-trial, e.g. the auditory go cue. The model makes choices by comparing *Q*_*L*_ and *Q*_*R*_ using a softmax choice rule. The model then compares the outcome of the choice with its corresponding reward prediction to calculate reward prediction errors (RPEs) used to update the weights *W*^*1*^ and *W*^*2*^. To have teaching signals proportional to our DLS DA signals, the model uses three different RPEs to update its parameters. For *W*^*1*^, the RPE is calculated using a ‘total’ reward prediction using all the inputs to the network (*Q*_*ch*_). For *W*^*2*^, the updates are separated into two pathways: the ‘stimulus’ and ‘constant’ pathway (Fig.3a Left, pink and aqua arrows respectively). The RPE in the ‘stimulus’ pathway is calculated using a ‘partial’ reward prediction 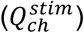 based on the stimulus inputs (*Vis stim. R* and *L*), whereas the RPE in the ‘constant’ pathway is calculated using a prediction based only on the constant input 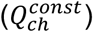. The stimulus-based RPE arising from this segregation is our account of DLS DA signals. This is motivated by the lack of DLS DA stimulus responses in trials with non-associated stimuli, despite high choice accuracy (Fig.2l,m). This learning rule yields updates that minimise three different losses through gradient descent: the ‘cortical’ loss equal to the total RPE^2^, and the two ‘cortical-striatal’ losses equal to the pathway-specific RPE^2^’s (see Methods). We term this model the ‘tutor-executor’ network because the ‘cortical’ learning (*W*^*1*^) *tutors* downstream ‘cortico-striatal’ learning (*W*^*2*^) by determining the relative salience of the inputs and balancing updates in the *executor* pathways to minimise its ‘total’ loss.

The model captured individual learning trajectories. Similar to mice, the model initiated learning by developing varying degrees of left/right bias (Fig.3b,c cf. Fig.1f,g). Subsequently, the model’s biases reversed as its psychometric slopes grew, exhibiting a similar diversity of left/right slope differences across trajectories to that seen in the mice (Fig.3d,e cf. Fig.1h,i). The model’s bias early in learning predicted bias and psychometric slopes later in learning, again showing striking similarity to mice data (Fig.3c,e cf. Fig.1g,i, Extended Data Fig.7a cf. Extended Data Fig.2g). As such, the model’s entire learning trajectories resembled those of mice, capturing their diversity and systematicity, and exhibiting similar sigmoidal accuracy curves over comparable time scales (Fig.3f-i cf. Fig.1j-m, Extended Data Fig.7c-e cf. Fig.1b-d, Extended Data Fig.7b cf. Extended Data Fig.1c). The simplicity of the model allowed us to derive expressions for its average learning dynamics which showed close proximity to our behavioural data and model simulations (Fig.3b,d,f-i, thick dashed lines). Therefore, a deep ‘tutor-executor’ RL network accounts for behavioural signatures of long-term learning within and across individuals.

DLS DA responses over learning were well captured by the model. We derived expressions for the trial-by-trial DA responses to stimuli and outcome using reward predictions from the ‘stimulus’ pathway (Methods). Similar to the empirical data, model-derived DA responses to stimuli grew over learning, reflecting stimulus-choice association (Fig.3j cf. Fig.2f, Extended Data Fig.7f cf. Fig.2d). The model-derived DA signals early in training predicted both the model’s slope difference (Fig.3k) and DA signals (Extended Data Fig.7g,h) late in training, similar to DLS DA data (Fig.2g, Extended Data Fig.4g,h). Thus, model-derived DA responses to stimuli across learning exhibited the diverse yet systematic progression of the empirical DA signals (Fig.3l-n cf. Fig.2h-j). Finally, model-derived DA responses to outcome also showed strong similarity to our data, encoding the difference between the value of each trial’s outcome and the reward predicted by the stimulus, i.e. RPEs in the ‘stimulus’ pathway (Fig.3o cf. Fig.2k, Extended Data Fig.7i-k cf. Extended Data Fig.4i, 6a,b). Taken together, the stimulus-based prediction errors of the deep ‘tutor-executor’ RL network mirror DLS DA signals throughout learning.

Analysis of the model average dynamics revealed a hierarchy of saddle points that explained the behavioural and neuronal trajectories. To find the average dynamics, we derived the average update to *W*^*1*^ and *W*^*2*^ across inputs and choices (Methods). Analysing the stationary points of the average dynamics (i.e., weight configurations where the average update goes to zero), revealed a series of saddle points. Saddle points have both stable and unstable manifolds, and in their vicinity learning momentarily slows down (36,37). These points span the entire learning process, starting from a ‘naïve’ weight configuration (*0*) and converging on a final expert global minimum (*4*) (Fig.3a Right). The saddle points establish a systematic flow through the parameter space, whereby the saddle points approached early in learning influence those approached later in learning.

Each stationary point in the model has a characteristic behavioural and dopaminergic signature, similar to mice behavioural strategies and DLS DA responses. Simulations start close to the first saddle point (*0* in Fig.3a Right) corresponding to a network configuration with all weights set to zero. Next, simulations that, for example, develop an initial right side bias learn in the direction of the *1R* saddle point, developing a strong association between the ‘constant’ input and *Q*_*R*_, evident in its corresponding weight configuration diagram (*1R* in Fig.3a Right). These simulations then move preferentially towards the next saddle point, *2R*, developing an association between the right stimulus and *Q*_*R*_ while maintaining a strong right bias. This happens as the simulations are still making mostly right choices (weight between ‘constant’ and *Q*_*R*_ is larger than between ‘constant’ and *Q*_*L*_) but start to learn the correlation between right stimuli and reward after right choices. Simulations approaching *2R* then move towards *3R*, where psychometric slopes emerge, and the bias starts to reverse. Here, the simulations maintain their association between the right stimulus and *Q*_*R*_ while developing a strong weight between ‘constant’ and *Q*_*L*_, all without using the left stimulus. Thus, in the vicinity of this saddle point the simulation infers correct left choices from the *absence* of the right stimulus, showing psychometric curves and DA signals similar to expert ‘right-associating’ mice. A mirror image of this trajectory is observed in simulations that initially develop a left bias (follow *1L, 2L* and *3L*), whereas more balanced simulations (those with a negligible initial bias) move from *1B* towards *4*, i.e., the global minimum. As such, the saddle points of the deep RL ‘tutor-executor’ network govern the learning trajectories (see saddle points visualised in Fig.3f-i and l-o), explaining their diverse yet systematic transitions between strategies and corresponding DA signals.

Certain features of our model are critical for capturing our data. The tutor-executor learning rule is not strictly necessary to capture our behavioural data; a deep RL model that minimises a single ‘total’ RPE through gradient descent achieves qualitatively similar results (Extended Data Fig.8 Left). However, such a single loss model uses teaching signals different from our recorded DLS DA signals (Extended Data Fig.8 Right). The depth of the network instead *is* critical in defining the learning trajectories; a shallow model does not reproduce the one-sided trajectories we observed empirically. In fact, depth is required to obtain a saddle point structure yielding the systematic transitions through strategies we observed in the mice (Extended Data Fig.9).

## Discussion

Our results show that in learning to make perceptual decisions from naïve to expert, mice display substantial individual variability while exhibiting systematic transitions through behavioural strategies. They initially increased their rate of reward harvest by making fast, non-accurate (biased) choices. This initial strategy determined their later strategy, i.e., which visual stimuli they use to solve the task. DA signals in the dorsolateral striatum (DLS) developed reflecting the stimulus-choice associations determined by each individual’s strategy. A deep reinforcement learning model trained using prediction errors analogous to empirical dopamine signals yielded diverse but systematic learning trajectories similar to those of mice. The learning trajectories were qualitatively governed by saddle points and their connecting manifolds, providing a formal account for how a biological learning mechanism can steer long-term learning.

Unlike conventional shaping methods for training animals that gradually change the task from easy to difficult, we chose to introduce mice to the full (i.e., relatively difficult) task from the onset of the experiment, and kept the task unchanged throughout learning. This training procedure allowed mice to freely explore and self-define their trajectories through the space of behaviour. In fact, the substantial early side biases, and later one-sided strategies we observed could be due to the higher starting difficulty of our task. The effect of task difficulty in developing biased strategies has been observed with other sensory modalities such as in whisker-based tasks (38). Future studies can examine the effects of various training curricula on learning trajectories. Our modelling framework allows for such investigation *in silico*, and for informing the design of curricula that accelerate learning.

Our work provides insights into the principles underlying the individual variability and systematicity of the mice’s learning trajectories. The saddle point structure emerging from the model’s architecture and learning rule reveals three main paths through the space of behaviour, giving a normative explanation for the diversity observed in mice. Within each path, the behaviour progressed systematically, such that the behaviour late in learning could be predicted by the behaviour in earlier stages of learning. The temporal resolution at which we observed systematic trajectories was of the order of tens of trials. However, there could also be behavioural strategy switches happening on faster time scales (39), which may be consistent with our observations. Further, in our model, the early variability in side bias emerges from imbalances in rewards after left and right choices. However, the early side bias in mice may also be influenced by other factors such as their position and handedness. As such, understanding the source of individual variability in these early days of learning requires further experiments.

DLS DA signals developed to encode stimulus-choice associations, such that stimuli whose presence – but not absence – informed choice and predicted reward, caused DLS DA release. Our model captures these signals using the stimulus-based prediction errors of its ‘stimulus pathway’. The model suggests that such local prediction errors could reflect the topography of connections between cortex and striatum, leading to different DA signals across striatal regions, as suggested by previous studies (40,24,41–43). For instance, our DLS DA signals were bilateral, in contrast to DA signals in more medial regions of dorsal striatum measured in a similar task (44). This difference could be due to DLS receiving more bilateral inputs from frontal and association cortical areas compared to dorsomedial striatum (45). The stimulus-based prediction errors observed here also differ from recordings of VTA DA neurons and their projections to ventral striatum in expert animals, which have been shown to integrate information across a wider variety of inputs and encode total, i.e. common currency, prediction errors, or even model-based prediction errors (46,47). The DLS DA signals we recorded modulate the strength of local cortico-striatal synapses (48). This, in turn, could rapidly regulate the size of stimulus-evoked responses in striatal neurons which form a functional closed loop by projecting back to midbrain DA neurons (49– 51), causing the changes in DA stimulus and outcome responses we observed throughout learning.

Signatures of stimulus-choice association emerged in RTs before being evident in choice accuracy. DLS DA signals reflected this early signature, growing in initial days when mice had a flat but biased psychometric curve and choices in trials with the stimulus on the biased side were becoming faster. Unlike in other studies (21,30), these DA signals were locked to stimulus onset and not choices. Nevertheless, these pre-outcome DA signals could contribute to reducing RTs by invigorating action (Extended Data Fig.5h) (27,52). Our DLS DA signals also differed from previously reported DA novelty signals (17,23,35) because they are absent in early days and grow over time. The absence of DA novelty signals could be because the visual stimuli we used were not salient enough in early days of learning, before their task-relevance was discovered. DLS DA stimulus-locked signals were similar in correct and error trials. Past studies measuring DA signals in the midbrain or ventral striatum of expert animals have shown smaller DA responses in error trials during perceptual decisions (52–54). This difference could be simply due to a difference in brain regions examined. Alternatively, this difference might emerge from different sources of errors: in our study errors reflect incomplete stimulus-choice association. However, in past studies of expert animals, errors are often caused by the limits of stimulus perception (55), leading to lower reward expectation in such trials.

We found that a hallmark of long-term learning to make perceptual decisions is the systematic progression through quasi-stages, connected by periods of more rapid change. Many other well-studied abilities–including face perception, semantic cognition, and numerical cognition–are characterised by similar structured transitions through strategies (56). Our model proposes a biological learning mechanism that exhibits such stage-like transitions, emerging from the hierarchy of saddle points that govern its learning dynamics. Interestingly, each saddle point corresponds to the network configuration that achieves the smallest loss using a subset of the available inputs or choices. Hence, the model explains the mice’s learning process as alternating between periods of discovering new inputs or choices, and periods of gradual association between the inputs and action value outputs (*Q*_*L*_ and *Q*_*R*_). The model also demonstrates that depth is a requirement of the circuit architecture, without which saddle points, and hence the characteristic learning stages, do not emerge. The same saddle point perspective has been offered as an explanation of stage-like transitions in semantic development (57). In contrast, ‘shallow’ reinforcement learning models commonly used in neuroscience can learn to perform the perceptual decision task (52), but do not capture the mouse learning trajectories.

Through the ‘tutor-executor’ learning rule, our neural network model reproduces the mouse learning trajectories with weight updates that are proportional to recorded DLS DA. A learning rule where different parts of the network minimise different loss functions, as used here, is unconventional in machine learning. However, this model accounts for both behavioural and DA data whereas standard gradient descent on a single total RPE^2^ only accounts for the behaviour. Thus, while behavioural signatures of learning are consistent with multiple closely related models, the DA signals favour the tutor-executor model. Future studies are needed to investigate whether the DLS DA signal causally drive long-term learning to make perceptual decisions.

An intriguing feature of the ‘tutor-executor’ learning dynamics is that, with extensive training, weight magnitudes transfer from *W*^*1*^ (‘cortical’) to *W*^*2*^ (‘cortico-striatal’) while maintaining the value of their product *W*^*2*^*W*^*1*^ (Extended Data Fig.10). This resembles the ‘transfer to striatum’ reported in past studies, where cortex was found to be crucial for task performance only in early stages of learning (58). Such a transfer can explain past results involving DLS in habitual behaviour (59), as the decrease in *W*^*1*^ weights leads to lower learning flexibility but could free the cortex for other tasks. This dynamic emerges from the difference in objective between *W*^*1*^ and *W*^*2*^ weights, since the total RPE teaching *W*^*1*^ reaches its minimum before the partial RPE’s in *W*^*2*^, causing *W*^*2*^ to keep growing and *W*^*1*^ to compensate by decreasing.

In summary, we have identified behavioural, dopaminergic, and computational principles underlying long-term learning trajectories. Given its widespread nature, a saddle point view of stage-like learning could generalise beyond our task, and could be a signature of learning in the ‘deep’ cortico-striatal circuit investigated in this study. Building on our results, future studies can investigate the dynamics of neural activity in other dopaminergic circuits or other brain regions during long-term learning.

## Acknowledgements

This work was supported by grants from the Wellcome Trust (213465/Z/18/Z) and ERC (funded by UKRI, EP/X026655/1) to A.L., grants from the Wellcome Trust (216386/Z/19/Z and 219627/Z/19/Z) and the Gatsby Charitable Foundation (GAT3755) to A.S., and grants from BBSRC (BB/S006338/1) and MRC (MC_UU_00003/1) to R.B. A.S. is a CIFAR Azrieli Global Scholar in the Learning in Machines & Brains program. M.F. is supported by a HFSP long-term postdoctoral fellowship.

## Data and Code Availability

The data generated in this project, as well as computer codes for data analyses and computational modelling will be shared publicly upon peer-review publication.

## Author Contributions

P.Z.H. and A.L. (Armin Lak) conceived and designed the experiment. P.Z.H., Ae.L. (Aeron Laffere), C.T., L.S., S.L.G. and A.L. performed the experiments. Y.L. shared viral constructs. S.L.G., Ae.L., P.Z.H., M.F., J.P., and A.L. analysed the data. S.L.G., R.B., A.L. and A.S. designed the model. S.L.G. and A.S. analysed the model. S.L.G., A.S. and A.L. wrote the manuscript with inputs from R.B., C.T. and M.F..

## Methods

### Mice

The data presented in this paper was collected from 28 male wild-type C57/BL6J mice, with their age ranging between 9 to 30 weeks. All experiments were conducted according to the UK Animals Scientific Procedures Act (1986) under appropriate project and personal licences.

### Surgical procedures

Animals were anaesthetised with isoflurane and were kept on a feedback-controlled heating pad (Stoelting 53810). Hair overlying the skull was shaved and the skin and muscles over the central part of the skull were removed. The skull was thoroughly washed with sterile saline. A head plate was attached to the bone posterior to bregma using dental cement (Super-Bond C&B). After the head plate fixation, we made craniotomies over the target areas and injected 300nl of AAV9-hsyn-DA2m into the right and/or left DLS (AP: +0.5 mm from bregma; ML: +/-2.5 mm from midline; DV: 2.8 mm from dura). This was followed by implantation of the optical fibre (core = 200 um, Neurophotometrics Ltd), which was secured to the head plate and skull using dental cement. Mice recovered for at least seven days following the surgery.

### Behavioural task

We trained mice in a complete psychometric visual decision making task from day 1 until expertise. Following surgery recovery, mice were first habituated to the experimenter for 2-3 days, followed by 2-3 days of habituation to the experimental rig and head-fixation. In each day of the experiment, mice were head-fixed with their body and hind-paws resting on a stable platform with a covering, and their forepaws resting on a steering wheel that could be rotated left and right. Each trial began after the wheel was held still for a short quiescence period (0.7-0.8s) (Extended Data Fig.1a). A sinusoidal grating stimulus of varying contrast (0%, 25% and 50%) was presented on either the left or right side of a screen in front of the mouse, followed by an auditory go cue 0.2s after visual stimulus onset. The go cue indicated the start of the interactive period, during which wheel movements were coupled with movement of the visual stimulus on the screen. The mouse was required to indicate the position of the stimulus by steering the wheel in the correct direction to move the stimulus to the centre of the screen, causing a water reward (3ul drop) to be delivered via a spout positioned close to the mouth. A variable ‘feedback delay’ (0.1-0.3s) separated the time of choice completion from reward delivery. Subsequently, the next trial started following an ‘inter-trial delay’ (2-3s). When an incorrect choice was made, the mouse was presented with a 0.5s auditory stimulus of white noise via speakers positioned near each ear and had a brief timeout period (2s) before the next trial. When a mouse responded incorrectly to an ‘easy’ high-contrast stimulus (50% contrast), there was a 50% chance that the same stimulus was repeated in the next trials, until the mouse responded correctly. A response window of 30s after the go cue was provided for the mice to make their choice. The behavioural experiments were delivered by custom-made software written in MATLAB (MathWorks) which is freely available (60).

### Imaging dopamine release

To measure dopamine release in the dorsolateral striatum (DLS), we employed fibre photometry. Photometry and behavioural data were collected simultaneously. We used chronically implanted optical fibres to deliver excitation light and collect emitted fluorescence (Neurophotometrics FP3002). We used multiple excitation wavelengths (470 and 415 nm), delivered on alternating frames (sampling rate of 40 Hz), serving as target and isosbestic control wavelengths, respectively.

The recorded photometry signal was pre-processed following steps described previously (61). We began by de-interleaving the recorded signal at 470 nm and 415 nm wavelengths. Both signals were then de-noised to remove short-pulse artefacts using a median filter with kernel size 5 (medfilt from scipy.signal). Subsequently, the signals were detrended with a zero-phase low-pass filter with a 10Hz cutoff frequency (2nd-order butterworth filtfilt from scipy.signal). Next, a photobleaching correction was applied to remove slow changes in the signal likely coming from fluorophore degradation due to light exposure throughout the recording session. To do this, we used a scipy filtfilt zero-phase high-pass filter with a cutoff frequency of 0.001Hz, thus removing signals varying with a timescale slower than 16 minutes. We then corrected for motion signals by fitting the 415 nm isosbestic to the 470 nm signal with a least squares polynomial fit of degree 1 (linregress, scipy.stats) and the resulting fitted signal was then subtracted from the 470 nm signal. Finally, this quantity (ΔF) was normalised through division by the baseline fluorescence (F, defined as a low-pass filtering of the denoised 470 nm signal with a cutoff frequency of 0.001Hz) to obtain ΔF/F which was subsequently z-scored per session to enable more accurate comparisons across days of recording.

High data quality was ensured by removing sessions with weak DA signals. We plotted the relative amplitude of the raw 470nm and 415nm signal per session. If this ratio was smaller than 1 the session was discarded, since in such sessions most of the pre-processed ΔF/F fluctuations came from variation in the isosbestic signal instead of the informative 470nm channel. We also discarded sessions where the maximum fluctuations were smaller than one standard deviation (i.e., Z<1).

### Video monitoring

The left eye was monitored with a camera (Teledyne Flir CM3-U3-13Y3M-CS) fitted with a zoom lens (Thorlabs MVL50M23) recording at 20 Hz. Front body movements were monitored with another camera (same model but different lens, Thorlabs MVL16M23) also recording at 20 Hz. Mice were illuminated with infrared light (850 nm, BW BWIR48) for the recording of eye and front body movements.

### Histology and fibre track quantifications

Histology after the experiment was performed to confirm successful fibre positioning. At the end of experiments, animals were deeply anaesthetised and perfused using 4% paraformaldehyde (PFA) and then decapitated. The brains were extracted, left in 4% PFA for 24h to post-fix in a refrigerator and then embedded in blocks of 1.5% agarose gel before collecting slices at 70 micrometre thickness using a vibratome (Leica VT1000 S). Slices were then stained with DAPI for 15 min (1:1000 solution), mounted onto glass, coverslipped, and imaged using an epifluorescence microscope (Leica).

### Behavioural data analyses

#### Behavioural data pre-processing

The behavioural data was pre-processed by removing the following trials: trials with response times more than 2 standard deviations above the mean per session, repeat trials (trials repeated after high contrast error trials) and trials where mice did not make a choice in less than 30s.

#### Behavioural metrics

The main behavioural metrics we used to analyse the mouse trajectories were accuracy, psychometric slope and bias. Accuracy was defined as the proportion of rewarded choices in all trials except for those without stimuli (i.e., zero-contrast trials), where choices were rewarded randomly. Psychometric curves were calculated per session, and plotted the proportion of rightward choices (i.e., P(‘Right’)) for different stimulus positions (left, right) and contrast values (0, 0.25, 0.5). The value of psychometric slope we used in all our analyses is that of the simplified psychometric curve collapsing across contrast levels to give left stim, zero-contrast and right-stim x-axis values. Left (right) slope was defined as the absolute difference in P(‘Right’) for left (right) stimulus and zero-contrast trials. Bias was defined as the difference between the P(‘Right’) on zero-contrast trials and 0.5, thus representing the imbalance of choices on zero-contrast trials in left and right directions.

#### Learning Trajectories

Each mouse was assigned a colour and cluster based on its learning trajectory. The rule for assigning colours used a weighted average of the difference between right and left per-session slopes over learning, where the weighting was equal to the sum of the left and right slopes on each session (Extended Data Fig.2c). The resulting average slope asymmetry metric was used to determine a colour for each mouse on a spectrum ranging from purple (for negative values) to orange (for values around 0) to green (for positive values).

The trajectories of the behavioural metrics were smoothed for better visualisation, highlighting their slow variation over learning. To do this, we used scikit-learn’s Gaussian process regression package (62) to fit a gaussian process with an RBF kernel (with tuneable scaling and length-scale) to the session-by-session metrics. The predict method of the fit gaussian process could then be used to estimate the smoothed value of the metric at different time points over learning.

A cluster label was assigned to each mouse to obtain cluster averages that highlighted the main trends in the diversity across learning trajectories. A dynamic time warping clustering algorithm was used to obtain these clusters (63). This algorithm first looks for the time warping that best clusters the trajectories by shape. The cluster centres are then computed as the barycenters with respect to the time warped mouse trajectories, yielding cluster centres that are similar in shape to the individual trajectories, thus solving the problem of averaging across mice that learn at different speeds. This clustering was applied to the smoothed right vs. left slope trajectories in Fig.1j and the resulting cluster labels were used to compute the cluster averages in all other behavioural and neural plots throughout the paper. The same colouring, smoothing, and clustering methods were applied to the model simulations to obtain plots similar to those produced for the experimental data.

#### Pupil analysis

We used DeepLabCut (64) to track several points on the mice’s left pupil throughout each task trial. We selected 4 points in the top, bottom, left and right portions of each mouse’s pupil and recorded the x and y coordinates of each point over time. For our pupil motion analysis (Extended Data Fig.2h), we defined the average x and y coordinate of these 4 points as the position of the pupil and investigated its horizontal (Δx) and vertical (Δy) motion. The average x and y coordinates were z-scored per session to enable more accurate comparisons across days. The alignment to stimulus onset was done as described in the neural analyses section. The analysis window used to compute pupil ‘stimulus responses’ was from 0.25-0.35s after the stimulus for the vertical motion and 0.5-0.6s for the horizontal motion.

### Neural data analyses

#### Event alignment and time warping

DLS DA recordings were aligned to task events that caused significant DLS dopamine release, as determined by a kernel regression (see below). These events were visual stimulus onset, choice completion (i.e., correct trials: visual stimulus arriving in the centre of the screen, and error trials: stimulus moving out of the screen) and trial outcome (reward/no reward). To do this, a fixed time period around each event (−0.5s to +1s) was selected and a fixed number of elements for the resulting aligned neural trace were chosen (i.e., 100). The DA recording was then linearly interpolated to obtain a value for each of the desired time points in the chosen time period (using scipy.interp1d). The average value in the time period before the event (−0.5-0s) was used as a baseline and subtracted from the event-aligned traces.

Time warping was used to visualise DA signals in a single continuous trace including all trial events. This was achieved by warping the DA signals such that a fixed number of time points represented the time course between each event. In this way, varying time periods between events were accounted for by allowing different time intervals between data points. We chose to have 40 time points before stimulus onset, 30 between stimulus onset and choice completion, 12 between choice completion and outcome and 60 post-outcome. The recorded DA signal was then interpolated to obtain the fluorescence values at the corresponding time points. This time warping allowed us to compute and visualise average DA signals across trials with varying durations.

#### Normalisation across sessions

In addition to the pre-processing steps described above, we further corrected for session-by-session variation in fluorescence levels by normalising the DA signals to the peak of the average DA response to reward delivery in zero-contrast (i.e., no visual stimulus) trials. To do this, we computed the average time warped DA trace per session, took the mean of the 3 largest values in the post-reward response (peak), and then subtracted the mean fluorescence levels 3 time points before and after the time of reward delivery (baseline) to obtain a single number (peak – baseline) that was used to divide the entire DA recording from that session by. This normalisation assumes that DA reward responses in zero-contrast trials should not vary session-by-session, as there are no cues that can predict the random reward delivery on these trials, meaning that the degree of ‘surprise’ (i.e., reward prediction error) should remain fairly constant throughout learning. This yielded normalised ΔF/F values that ranged from 0-1, expressing the signal as a proportion of the zero-contrast reward response in each session, thus resulting in more accurate averages across sessions and mice and more interpretable fluorescence values that could be compared with the model-derived DA signals.

#### Analysis windows

We defined average DA responses to 3 events (stimulus onset, choice completion and reward) which we use in several analyses throughout the paper. These were defined as the average DA signal in a specific time window after each event, relative to the signal before the event.

For the ‘stimulus response’, we defined an analysis window from +0.2s to +0.35s post-stimulus onset and took the peak response of a moving average (window size = 10) of the signal in that time window. We then subtracted a baseline fluorescence value (defined as the mean ΔF/F in a window −0.1s to +0.1s around stimulus onset) from the peak estimate to obtain the ‘stimulus response’ on each trial.

For the DA response to choice completion (‘stim. centre’ on correct trials), we used the time warped traces to define the analysis windows for the peak and baseline estimates. We did this because the times between stimulus, choice completion and reward delivery/absence are variable and can be short, so using time warping is an accurate method to obtain isolated estimates of DA release values uniquely around this event. To do this, we used the entire 12 elements of the time warped trace between choice complete and reward delivery/absence and found the peak value of its moving average (window size = 6). From this, we subtracted the mean of the time warped DA signal one time point before and after the choice complete event.

For the DA response to reward delivery/absence we used the same procedure as for the ‘stimulus response’. We only changed the analysis window from which we calculated the peak response, i.e., 0-1s after the reward time. The baseline was similarly defined using the average signal in a window −0.1s to +0.1s around the event.

For most neural analyses we defined a combined ‘outcome response’ which is the addition of DA responses to choice completion and reward delivery/absence. This was done because in rewarded trials, the ‘stim. centre’ cue is a perfect predictor of upcoming reward and thus rapidly acquires value, causing the DA responses to reward to become smaller. We did not want to include this learning process in our analyses as it does not involve decision making (i.e., is Pavlovian). Hence, by adding the responses for the two events, we obtained an outcome signal that did not change until events before choice completion became associated with reward.

#### Kernel regression

To find the events that caused significant variation in DLS DA release levels, we used a kernel regression algorithm. This algorithm works by regressing binary variables indicating the time period around events onto full trial DA signals. The design matrix (*X*) has a column per time point around the events of interest (regressors) and a row per time point of the DA signal, which was concatenated for all the trials used in the regression to form the dependent variable (*y*). The algorithm then finds the optimal scaling (*β\**) of the regressors in *X* which produces a prediction of the concatenated DA signal (*ŷ\**) that minimises the mean squared error Σ_i_ *(y*_*i*_ *– ŷ*_*i*_)^*2*^,

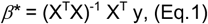

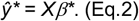

The resulting elements in *β\** compose the ‘kernels’ for each event. The benefits of using kernel regression over an event-aligned average is that it isolates the effect of each event on the DA signal, removing the influence of other events occurring shortly before or after. This isolation is achieved as long as there is enough jitter between events across trials; if two events are separated by a fixed delay in all trials the regression will not find isolated kernels due to correlated regressors.

To account for changing DA signals over learning in our kernel regression, we split all trials into 4 bins with increasing psychometric slope. We also performed separate regressions for rewarded and unrewarded trials. The events we considered were stimulus onset, choice start, choice completion and reward delivery/absence, which resulted in a total of 32 kernels (Extended Data Fig.3b). The explained variance for each kernel was computed by comparing the 5-fold cross-validated R^2^ of a ‘full’ model with all the kernels against that of a model without the kernel being assessed.

We used a custom implementation of this algorithm written in Python, which can be found in our analysis code repository.

### Deep linear RL model

#### Architecture

The model is a 3-layer deep neural network with linear activation functions (Fig.3a Left). We denote the weight matrix connecting the input layer to the hidden layer *W*^*1*^, and the matrix connecting the hidden layer to the output layer *W*^*2*^, such that the function computed by the network is given by

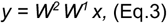

where *x* is the vector of inputs and *y* is the vector of outputs. The weights in both *W*^*1*^ and *W*^*2*^ are constrained to be nonnegative, and *W*^*1*^ is constrained to be a diagonal matrix. These constraints were chosen for simplicity and are not strictly necessary to capture the data. The network has two binary input neurons which encode the presence or absence of the right and left visual stimulus and one input neuron that has an activation of 1 for every trial, representing any non-stimulus cues that the mice may use to make choices on each trial, e.g. the auditory go cue. Thus, the network receives three different input vectors depending on the trial type,

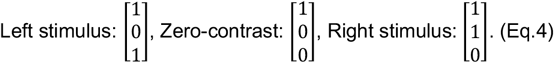

We opted not to model the different contrast levels on different trials (25% and 50%) because in most mice we did not see a significant difference in accuracy between these two levels.

In every trial, the input vectors are multiplied by the weights in *W*^*1*^ and *W*^*2*^ to obtain the activation of the two output neurons which encode the learned value of left and right choices, *Q*_*L*_ and *Q*_*R*_. These action values are used to determine choice through a softmax function with inverse temperature *β* that determines the choice probabilities,

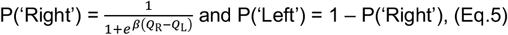

from which a choice is sampled on each trial.

For each simulation, the initialisation of *W*^*1*^ was sampled from a gaussian distribution centred on fit values of the initial stimulus input weights (identical for left/right) and the constant input weight. Similarly, the initialisation of *W*^*2*^ was sampled from a gaussian distribution centred on fit values of the stimulus pathway weights (Fig.3a Left, pink – all identical) and the constant pathway weights (Fig.3a Left, aqua – all identical). The softmax inverse temperature parameter *β* and learning rate *α* were also sampled from a gaussian centred on a value fit to the experimental data. The fitting procedure is described in the corresponding subsection. Simulations were run for 10,000 iterations (i.e., trials) and those that reached 70% accuracy in less than 8,500 iterations (approx. highest number of trials required for mice to learn) were included in our analyses.

#### Learning rule

##### a. ’Tutor-executor’ gradient descent

We refer to the model presented in Fig.3 as the ‘tutor-executor’ model due to its learning rule, which uses different reward prediction errors (RPEs) to train the weights in *W*^*1*^ and *W*^*2*^. The updates minimise three different losses through stochastic gradient descent (SGD): the ‘cortical’ loss for the weights in *W*^*1*^, the ‘stimulus cortico-striatal’ loss for the weights in the stimulus pathway of *W*^*2*^ (Fig 3a Left, pink), and the ‘constant cortico-striatal’ loss for the weights in the constant pathway of *W*^*2*^ (Fig 3a Left, aqua). Each of these losses is a RPE^2^ comparing predictions based on different subsets of inputs against trial outcome,

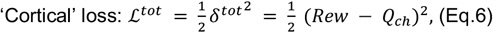

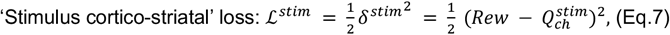

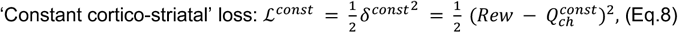

where *Rew* is a binary variable indicating whether the trial was rewarded or not, and the subscript *ch* indicates the choice made on each trial (left/right). Here *Q*_*ch*_ is the ‘total’ *Q*-value that uses all inputs to form its predictions, while 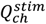 and 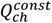 are ‘partial’ *Q*-values based on the stimulus and constant inputs in turn,

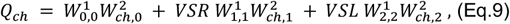

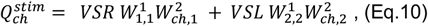

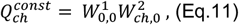

where *VSR* and *VSL* are the binary inputs indicating the presence or absence of the right and left stimulus respectively and *ch* used as a subscript for the weights is 0 for left and 1 for right choices.

Gradient descent on these losses yields updates which depend on the trial outcome and choice,

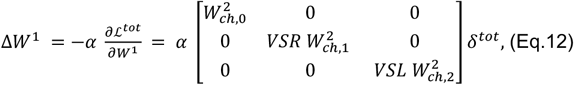

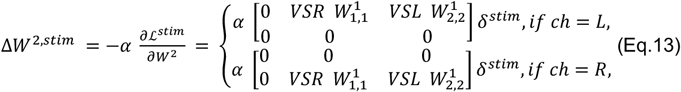

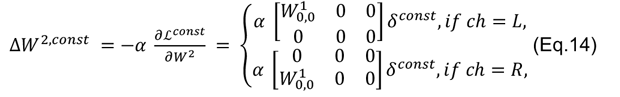

where *α* is the learning rate and Δ*W*^2^ = Δ*W*^2,*stim*^ + Δ*W*^2,*const*^. Notice how the updates for *W*^*1*^ are proportional to the total RPE, *δ*^*tot*^; the updates for the stimulus pathway in *W*^*2*^ are proportional to the stimulus-based RPE, *δ*^*stim*^; and lastly, the updates for *W*^*2*^’s constant pathway are proportional to the constant-based RPE, *δ*^*const*^. Interestingly, the general tendency of the learning rule is to minimise the total ‘cortical’ loss, ℒ^*tot*^, as the learning in *W*^*1*^ *tutors* downstream learning in *W*^*2*^ by determining the relative salience of the inputs and balancing updates in the *executor* pathways.

##### b. Single-loss gradient descent

This learning rule updates *W*^*1*^ and *W*^*2*^ through stochastic gradient descent (SGD) to minimise a single total RPE^2^. This corresponds to the conventional method of training deep RL networks, where all parameters are updated to minimise a single loss and thus share the same objective (65). The loss we use for this learning rule is the same as the ‘cortical’ loss in the tutor-executor model,

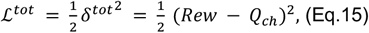

where *Rew* is a binary variable indicating the outcome of a particular trial and *Q*_*ch*_ is a reward prediction calculated based on all inputs to the network. The updates for *W*^*1*^ and *W*^*2*^ can be written as follows

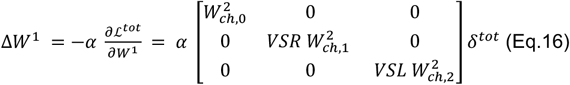

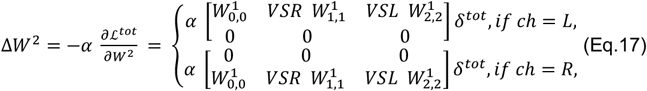

where *α* is the learning rate and *VSR* and *VSL* are binary variables indicating the presence or absence of the right and left stimulus respectively. Here, all the updates are proportional to the same total RPE,*δ*^tot^.

#### Model-derived dopamine signals

Our neural network model captures empirical dorsolateral striatal dopamine (DLS DA) signals through the weights in its stimulus pathway. Over learning, the model reproduces DLS DA outcome responses with a stimulus-based reward prediction error (RPE),

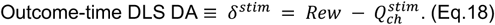

This differs from conventional temporal difference reward prediction errors (TD-RPEs) commonly used in DA studies in that it does not use the full reward prediction based on all inputs to define the RPE, instead comparing trial outcome with a prediction based only on the stimulus inputs. This was motivated by our matched accuracy analysis in Fig.2l,m, which showed that DLS DA at both stimulus and outcome time does not reflect reward predictions based on the constant input (i.e., the absence of a stimulus).

At stimulus time, in accordance with the TD-RPE hypothesis (15), the model captured DLS DA signals with a pre-choice reward prediction. Importantly, this prediction is also based only on the stimulus inputs,

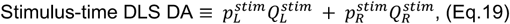

where 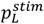 and 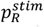 are the left and right choice probabilities determined by the stimulus-based *Q*-values passed through the choice function in Equation 5. Here, there is no need to subtract the value of the previous state as there is no cue before the stimulus that is predictive of reward.

To test whether the network could be trained with a learning signal analogous to our DLS DA recordings, the tutor-executor model uses a learning rule based on these ‘partial’ prediction errors to update its weights. Specifically, the updates of the *W*^*2*^ stimulus pathway weights are proportional to the stimulus-based RPE, *δ*^*stim*^ (Equation 13). Comparing the evolution of *δ*^*stim*^ with the DLS DA outcome response over learning shows a striking similarity (Fig.3o, Extended Data Fig.7f,i-k), suggesting that a similar dopamine-based learning mechanism could be governing the learning process of the mice.

#### Average dynamics

Deriving the average dynamics of the model allowed us to obtain an analytical description of its learning process. We took the continuous time limit of the average gradient descent updates for *W*^*1*^ and *W*^*2*^ from the tutor-executor and single-loss learning rules, averaging over trial type (left stim., right stim., or zero-contrast) and choice (left or right). This yielded a 9-dimensional system of coupled differential equations describing the evolution of each weight in the network. For sufficiently small learning rates *α*, this ‘gradient flow’ limit provides a good description of the average dynamics for both learning rules. The resulting differential equations were numerically integrated to obtain average weight trajectories over training time. To capture the three main types of learning trajectory (i.e., left-associating, balanced and right-associating), we initialised the integration with a network configuration yielding different degrees of initial choice bias (i.e., imbalanced connections from const. to *Q*_*L*_ and *Q*_*R*_). We then overlayed the resulting trajectories on those from trial-by-trial simulation (thick dashed lines in model figures).

##### a. Tutor-executor average dynamics

The average dynamics of the network weights for the tutor-executor learning rule are governed by three main differential equations; one for each of the cortical, stimulus cortico-striatal and constant cortico-striatal weight subsets (black, pink, and aqua in Fig.3a Left). These can be derived by taking the average over trial-types and choices of the gradient descent updates in Equations 12-14. Doing this for the cortical weights in *W*^*1*^ we obtain

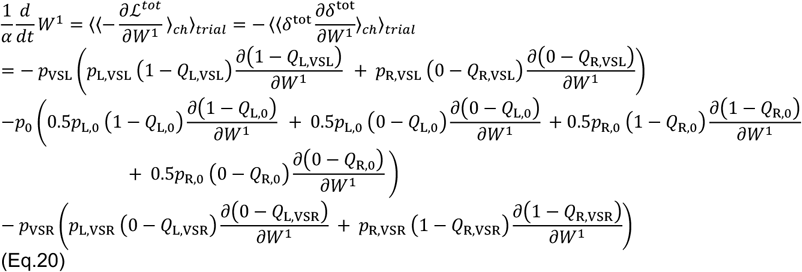

where *t* is a continuous time variable counting the number of trials; *δ*^*tot*^ represents the total RPE; *p*_VSL_ = *p*_VSR_ = 0.45 and *p*_0_ = 0.1 are the probabilities of there being a left vis. stim., right vis. stim., and zero-contrast trial; and lastly *p*_A,B_ and *Q*_A,B_ indicate the choice probabilities and total *Q*-values for choice A in a trial of type B. The choice probabilities are calculated using the sigmoidal choice rule in Equation 5. The partial derivatives of the *Q*-values can then be expanded by writing them in terms of the network weights (Equations 9-11) and differentiating w.r.t. the *W*^*1*^ matrix.

The same procedure can be followed to find the differential equations for the weights in the stimulus and constant pathways of *W*^*2*^, with gradient flow equations

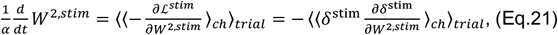

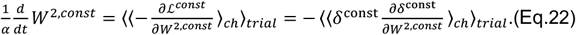

##### b. Single-loss average dynamics

The single-loss average dynamics can be derived in a similar fashion, where now all the weights are minimising the same loss function (Equation 15), yielding the following gradient flow equations

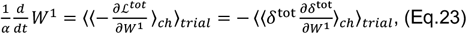

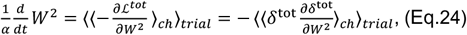

which can be expanded as exemplified with the tutor-executor dynamics.

##### Saddle Points

Saddle points in the model’s learning dynamics provide an explanation for the systematic transitions between behavioural strategies and dopamine release patterns observed in the mice. We used the average dynamics derived in the sections above to demonstrate the existence of these saddle points. We first made informed guesses at stationary points by looking for network configurations where the average dynamics go to 0, and then investigated the dynamics around these points to verify their nature. These derivations can be found in the following Mathematica notebooks:

https://www.wolframcloud.com/obj/samuelliebanagarcia/Published/tutor_executor_stationary_points.nb

https://www.wolframcloud.com/obj/samuelliebanagarcia/Published/single_loss_stationary_points.nb

We also provide evidence for heteroclinic orbits connecting the saddle points, represented by the arrows in Fig.3a and Extended Data Fig.8a. To do this, we used the string method (66) to find the minimal energy path between each pair of saddle points. This allowed us to distinguish between points that are directly connected by such paths (i.e., heteroclinic orbits), and points that are only connected through another one of the saddle points. The orbits discovered by the string method are shown in the Mathematica notebooks, and their schematic form is shown in Fig.3a and Extended Data Fig.8a.

#### Fitting procedure

To reproduce the learning trajectories in the data with our model, we fit the value of the network weights at initialisation and the *β* parameter of the choice function. Specifically, we fit the initial *W*^*1*^ weights for the constant and stimulus inputs (equal for left/right), and the initial *W*^*2*^ weights for the constant and stimulus pathways (all equal within each pathway). We took advantage of our expressions for the average dynamics of both the tutor-executor and single-loss learning rules, and minimised the mean squared error between the three trajectories emerging from integrating the dynamics starting from a small amount of left, right and no bias, and the three trajectory clusters from the data (derived from Fig.1j). The parameters were fit using momentum-based gradient descent by comparing the learning trajectories from the data shown in Fig.1j-m and Fig.2d with the same trajectories derived from integrating the average dynamics equations. This proved much faster than simulation-based methods. Lastly, the learning rate *α* was fine-tuned by hand to obtain learning trajectories that were stable and learnt in a similar number of iterations (i.e., trials) as the mice. The resulting parameter values were used as the means of gaussians from which the parameters of each simulated network were sampled for the simulations shown in the figures across the paper.

### Software packages

All data analyses were performed using custom code written in Python 3 using standard analysis and plotting libraries: numpy, scipy, matplotlib and seaborn. For the model, the JIT compilation and automatic differentiation capabilities of JAX were used to accelerate and simplify gradient calculations.

## Extended Data Figures

**Extended Data Fig.1.**
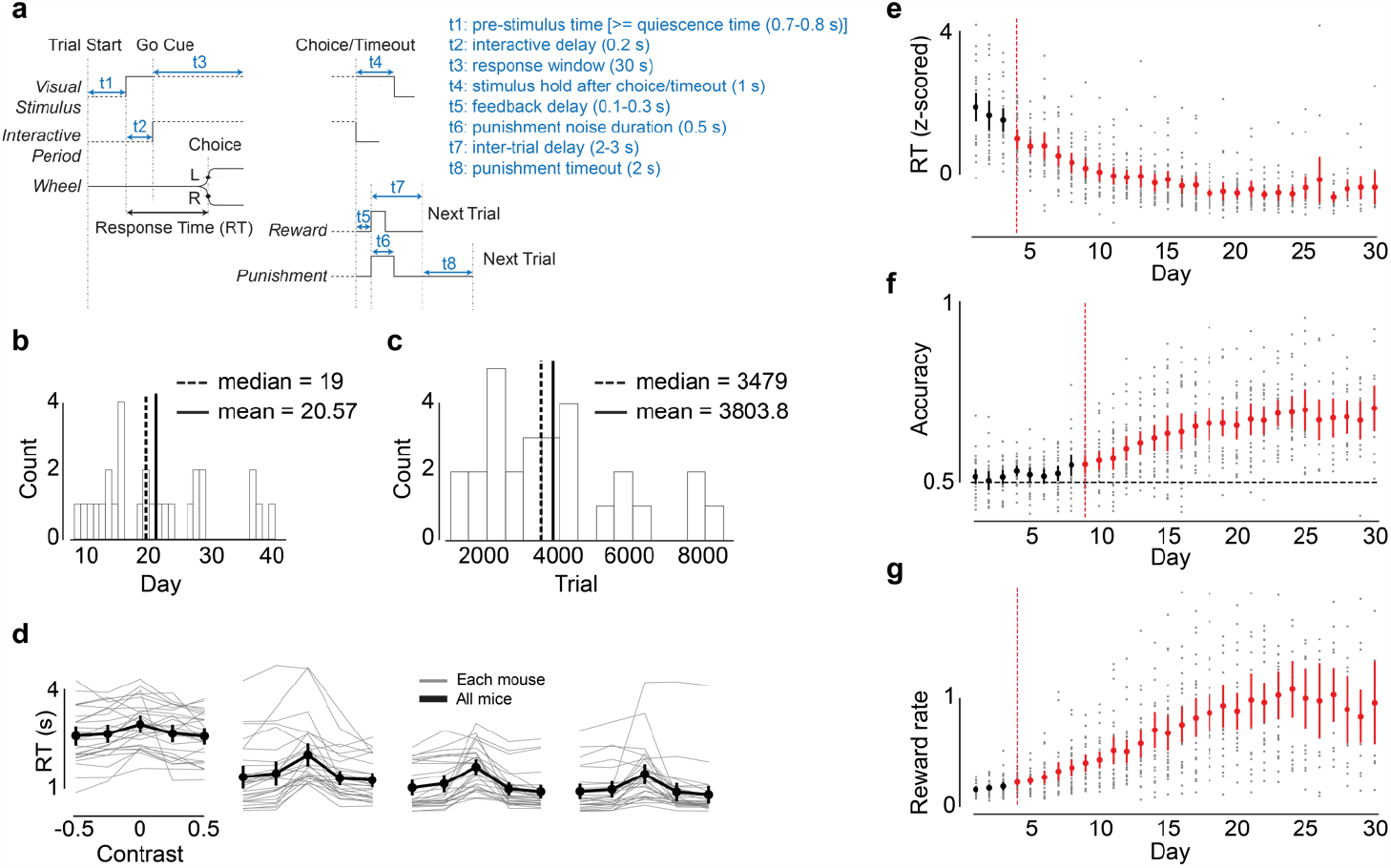
Learning a visual decision task from naïve to expert. **a**, The temporal structure of the task within a trial. Words in regular font indicate trial events, words in italics are labels for the traces in the timing diagram, and the solid vs dashed style of the traces indicate fixed vs variable time periods respectively. **b**, Histogram of the number of days that mice required to reach 70% accuracy. **c**, Histogram of the number of trials that mice required to reach 70% accuracy. **d**, Chronometric curve over quartiles per mouse (grey) and averaged across all mice (black). Negative (positive) contrast values indicate stimuli presented on the left (right) side of the screen. RT (s) indicates the mean response time (time from stimulus onset to choice completion) per contrast value in seconds, averaged first across trials in each session, and then across sessions in each quartile. Quartiles are defined per mouse by dividing days into 4 groups, with any remainder added to the last group. Error bars indicate the 95% confidence interval across mice. **e**, Z-scored RTs over training days per mouse (grey) and averaged across all mice (black/red). Red dots indicate data points significantly different from day 1 averages (*p*<0.05, estimated via two-sided t-test). Z-scoring was performed by standardising each mouse’s mean RT per day using data from all its days. **f**, Accuracy over training days. **g**, Reward rate over training days, calculated as the choice accuracy divided by the mean (not z-scored) RT for each day.

**Extended Data Fig.2.**
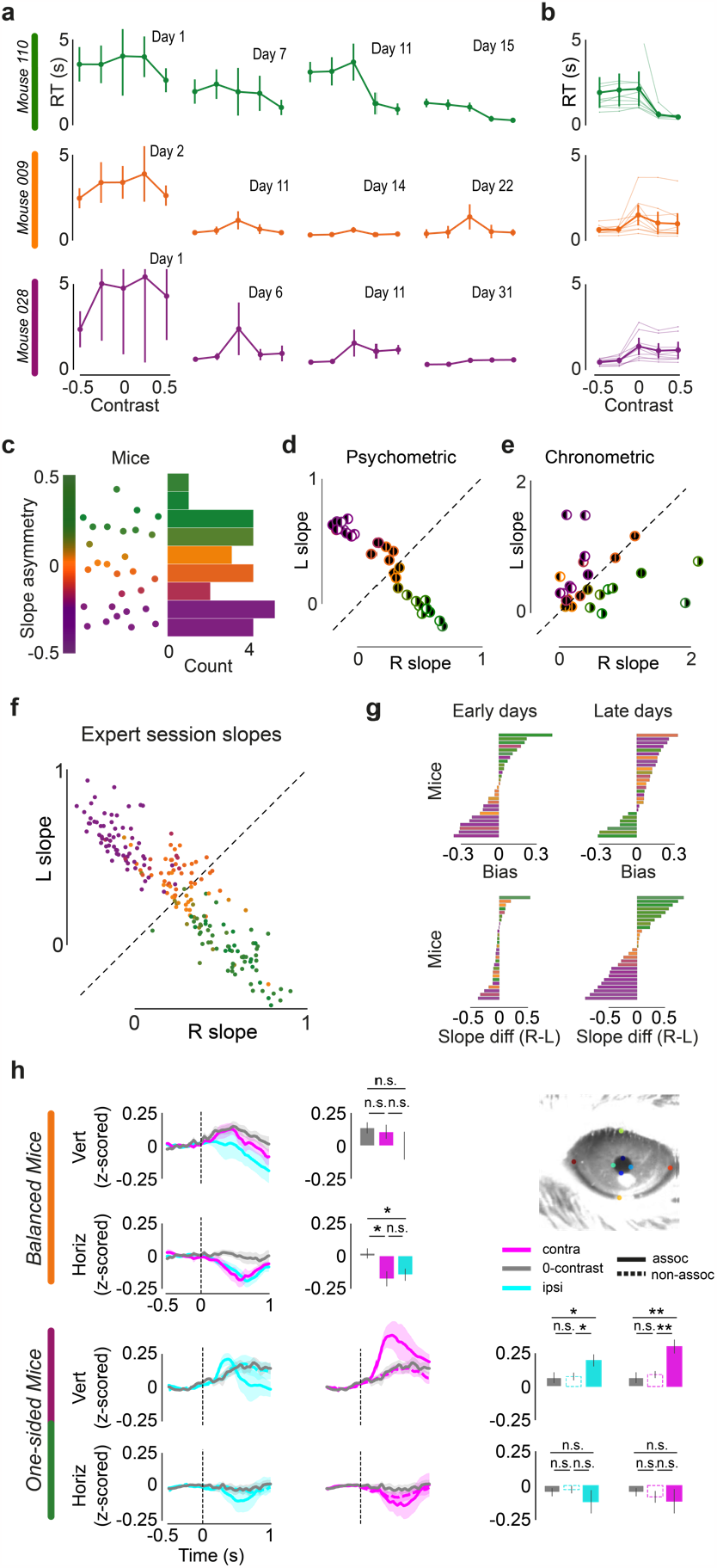
Learning from naïve to expert across mice. **a**, Chronometric curves for the same days and mice as in Fig.1d. Error bars indicate 95% confidence interval across trials in each day. **b**, Expert session chronometric curves per mouse (thin) and averaged per trajectory cluster (thick): right-associating (green), balanced (orange) and left-associating (purple). Cluster labels for each mouse obtained from Fig.1j. Error bars indicate 95% confidence interval of the average across mice in each cluster. **c**, Colour scheme (left), scatter (middle), and histogram (right) of average psychometric slope asymmetry per mouse. Slope asymmetry was calculated as 0.5 times the weighted average of the difference in R-L slopes over all sessions in a mouse’s learning trajectory. The weightings in the average are the sum of both R+L slopes per session. **d**, Scatter of average right vs. left expert session psychometric slopes for each mouse. Stroke colour indicates the average slope asymmetry of each mouse. The fill of the left/right half of each circle (black/white) indicates the statistical significance of the left/right slopes (black means significant). **e**, Scatter of average right vs. left expert session chronometric slopes for each mouse. Chronometric slopes were defined as the difference between the median RT on zero-contrast trials and trials with right/left stimuli. **f**, Scatter plot of expert session right vs. left slopes, coloured by the average slope asymmetry of the mouse trajectory they belong to. **g**, Bar plots showing the average bias and slope difference in early (4-8) and late (final 5) days for each mouse. Colours come from average psychometric slope asymmetry (see panel c). **h**, Average vertical and horizontal eye movements (pupil motion) aligned to stimulus onset in balanced and one-sided mice for stimuli contralateral (pink) and ipsilateral (blue) to the pupil, as well as on zero-contrast trials (grey). In one-sided mice, solid lines indicate pupil motion in response to ‘associated’ stimuli and dashed lines to ‘non-associated’ stimuli. Bar plots quantify pupil motion by taking the average in analysis windows from 0.25-0.35s after stimulus onset for the vertical motion and 0.5-0.6s for horizontal motion. * and ** indicate *p* < 0.05, *p* < 0.005.

**Extended Data Fig.3.**
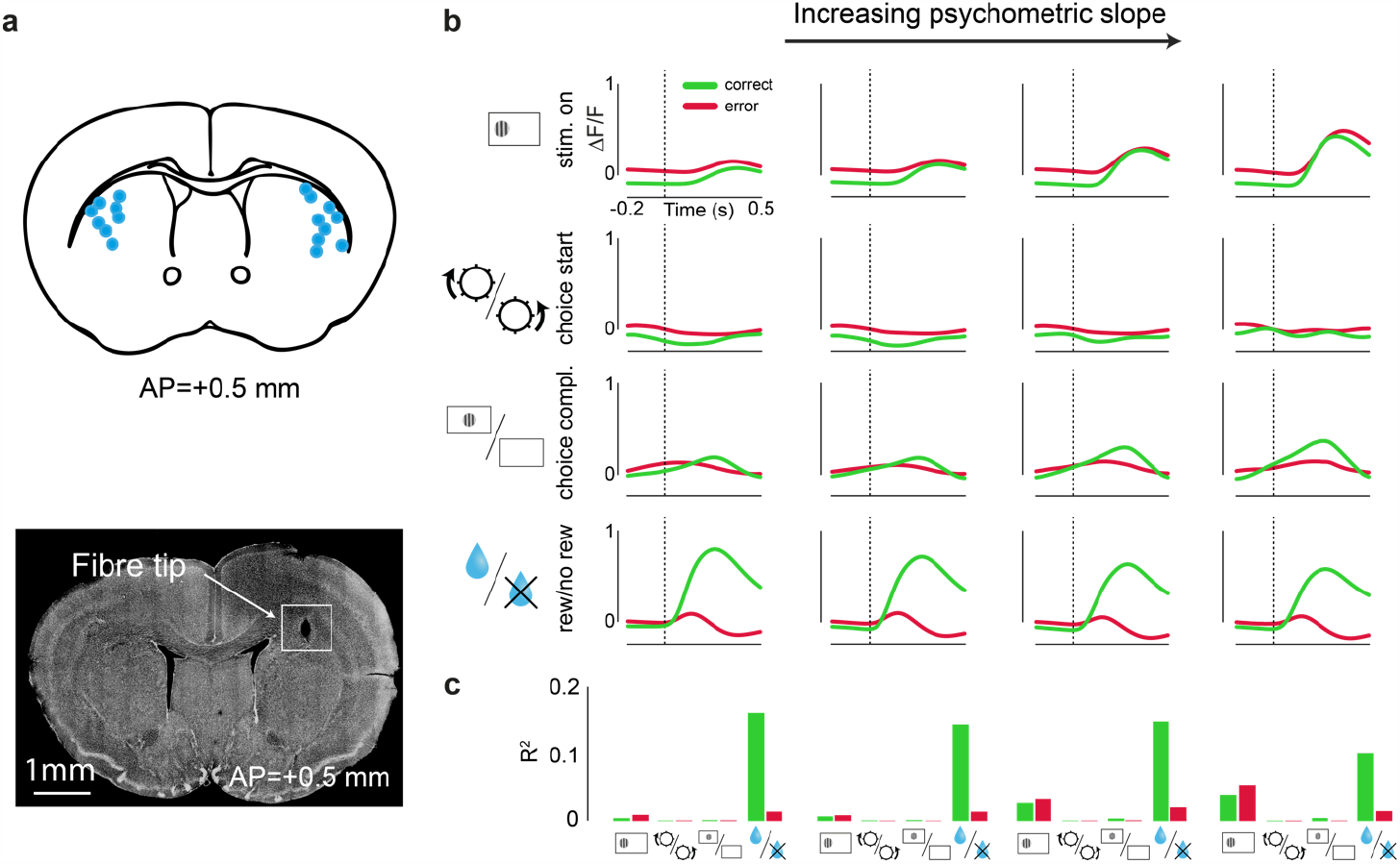
Dopamine signals in dorsolateral striatum during task learning. **a**, Top, the locations of the optic fibre estimated from post-mortem histological examinations overlaid on brain slice schematic. Bottom, example brain slice showing fibre tip. **b**, Regression coefficients of a kernel regression of DLS DA signals on different events in a trial: stimulus onset, choice start, choice completion, and outcome (reward/no reward). The regression uses the time points around each event as categorical regressors, and then finds the least squares regression coefficients predicting the original signal. The data was divided into correct and error trials, and was grouped into four bins of increasing psychometric slope. The kernel regression was performed independently for each subset of data. **c**, R^2^ for each trial event used in the kernel regression, calculated independently for correct and error trials and for each level of psychometric slope.

**Extended Data Fig.4.**
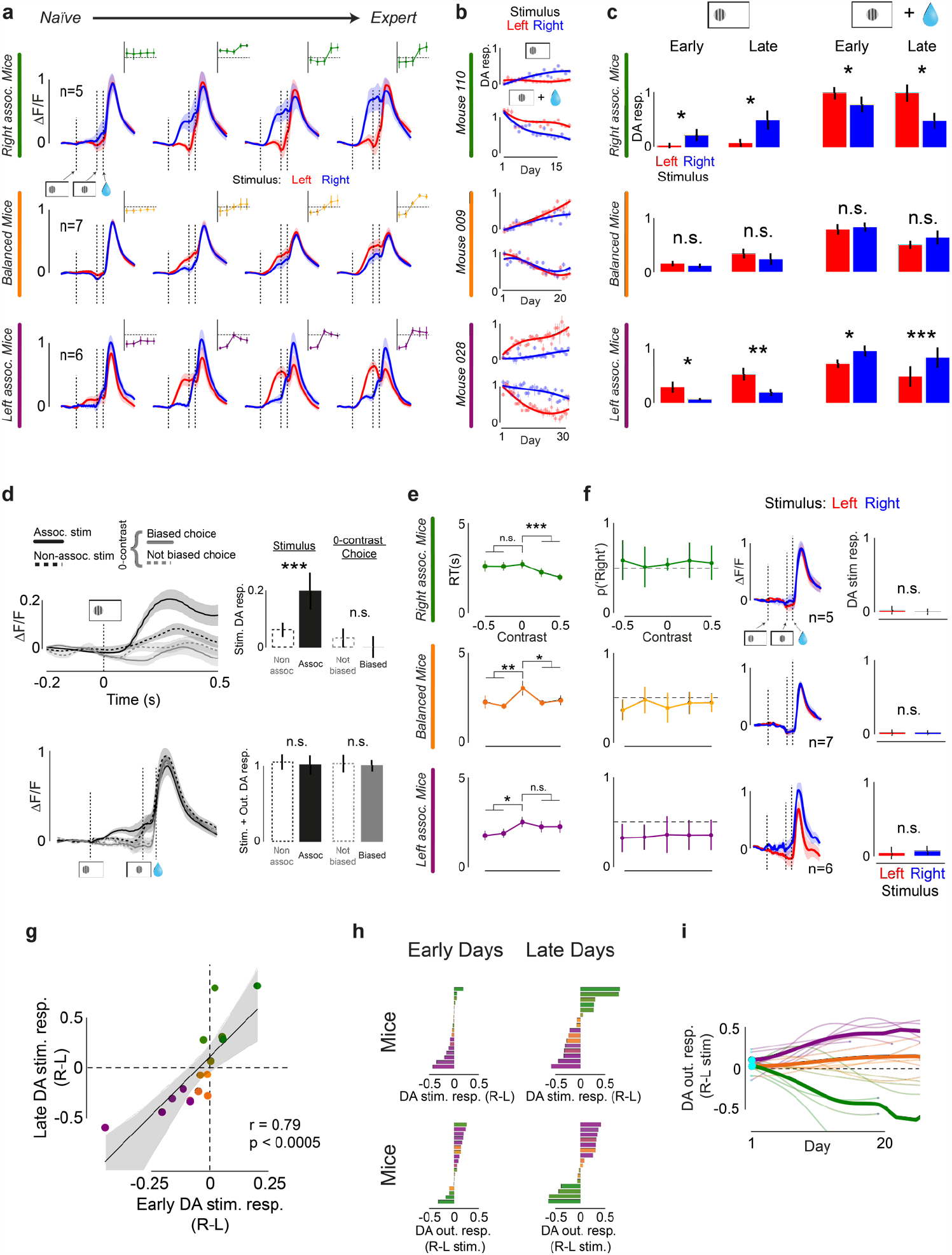
Dopamine signals during learning across mice. **a**, Average time warped DLS DA signals across mice in each cluster, plotted over quartiles for correct trials with stimulus on the left (red) and right (blue), c.f. Fig.2c. Vertical dashed lines indicate stim. onset, stim. centre, and reward delivery time. Insets show average psychometric curves across mice in each cluster for every quartile. Error bars in both neural and behavioural plots indicate 95% confidence interval of average across mice. **b**, Average stimulus and outcome DA responses over days in correct trials with stimulus on the left (red) and right (blue) for the three example mice from Fig.1d and Fig.2c. Error bars indicate +/-s.e.m. across trials. Data points fit with a 3rd degree spline to visualise trend (scipy.interpolate.UnivariateSpline), c.f. Fig.2d. **c**, Quantification of DA signals from Fig.2e. Left, bar plots showing average DA stimulus responses across mice in each cluster for correct trials with stimulus on the left (red) and right (blue) in early days (accuracy n.s. greater than 0.5) and late days (accuracy n.s. smaller than 0.7). Error bars indicate 95% confidence interval of average across mice. *p*-values calculated using two-sided t-test. Right, same as left but for outcome responses. **d**, Top left, average early days (accuracy n.s. greater than 0.5) stimulus-aligned DA signals in correct trials for one-sided animals. Solid and dashed black lines show signals in trials with associated and non-associated stimuli respectively. Solid and dashed grey lines show DA signals on zero-contrast trials (aligned to the time when the stimulus would have been presented, i.e., 0.2 s before the auditory go cue) for choices in and opposite to the direction of each mouse’s early bias. Bottom left, average early day time warped DA signals in correct trials. Top right, average early day DA stimulus responses in correct trials using the same legend as in top left. Bottom right, the sum of average early day DA responses to stimulus and outcome. *p*-values calculated using two-sided t-test. Error bars for stim.-aligned and time warped plots indicate +/-s.e.m., and bar plot error bars indicate 95% confidence interval, both across mice. **e**, Early day (accuracy n.s. greater than 0.5) chronometric curves for the three clusters of mice. Error bars indicate +/-s.e.m. across mice. *p*-values calculated using two-sided t-test. *, ** and *** indicate *p* < 0.05, *p* < 0.005 and *p* < 0.0005. **f**, Left, first/second day of training average psychometric curves for the three clusters of mice. Error bars indicate 95% confidence interval of average across mice. Middle, first/second day of training time warped DA signals in correct trials. Error bars indicate +/-s.e.m. across mice. Right, first/second day of training DA stimulus responses in correct trials. Error bars indicate 95% confidence interval of average across mice. **g**, Regression of early difference in DA responses to R-L stimuli (average across days 4-8) against late difference in DA responses (average across final 5 days). Each point represents a mouse. *p*-value calculated from the exact distribution of r. **h**, Average difference in DA responses to right and left stimuli (R-L) and difference in DA rewarded outcome responses after R-L stimuli in early days (4-8) and late days (final 5) for each mouse, c.f. Extended Data Fig.2g for behavioural data. Colours come from average psychometric slope asymmetry (see Extended Data Fig.2c). **i**, Difference in DA responses to rewarded outcomes after R-L stimuli over days per mouse (thin) and for the 3 clusters from Fig.1j (thick). Number of days limited to 25 for better visualisation, c.f. Fig.2f.

**Extended Data Fig.5.**
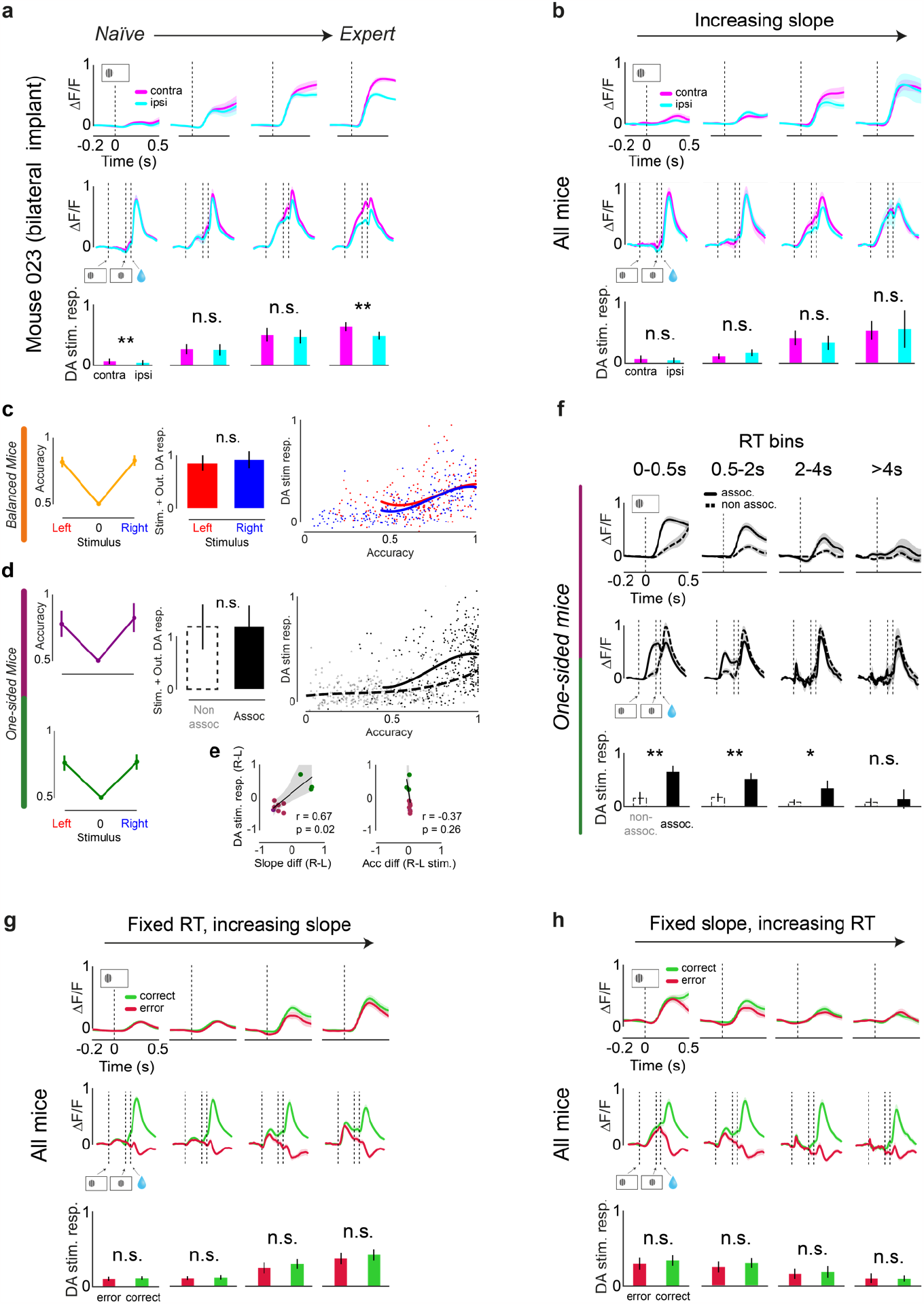
Dopamine signals during learning split by stimulus laterality, response times and trial outcome. **a**, Average DA stimulus-aligned signals (top), time warped signals (middle) and bar plots with average DA stimulus responses (bottom) for rewarded trials from each quartile of an example mouse with bilateral implants split by stimulus laterality w.r.t. implanted fibre. Error bars indicate +/-s.e.m. (top, middle) and 95% confidence interval (bottom) across days. *p*-value calculated from two-sided t-test. **b**, Average DA stimulus-aligned signals (top), time warped signals (middle) and bar plots with average DA stimulus responses (bottom) in rewarded trials for increasing psychometric slope values of all mice split by stimulus laterality with respect to the implanted fibre. Error bars indicate +/-s.e.m. (top, middle) and 95% confidence interval (bottom) across mice. *p*-value calculated from two-sided t-test. **c**, Left, balanced mice average accuracy in days with matched choice accuracy for trials with left (red) and right (blue) stimulus. Error bars indicate 95% confidence interval of average across days. Middle, balanced mice bar plots showing sum of average DA stimulus and rewarded outcome responses in trials with left and right stimuli in matched accuracy sessions. Error bars indicate 95% confidence interval of average across days. Right, balanced mice scatter plot of average DA stimulus responses to right (blue) and left (red) stimuli against accuracy. Each point represents a day. Data points fit with a 3rd degree spline to visualise trend (scipy.interpolate.UnivariateSpline). **d**, Same plots as in c reproduced for one-sided mice. **e**, Left, regression of difference in R-L psychometric slopes against difference in DA responses to R-L stimuli for matched accuracy sessions from one-sided mice. Right, regression of difference in choice accuracy in R-L stimulus trials against difference in DA responses to R-L stimuli for matched accuracy sessions from one-sided mice. Each point represents a recorded hemisphere. *p*-value calculated from the exact distribution of r. **f**, Average DA stimulus-aligned signals (top), time warped signals (middle) and bar plots with average DA stimulus responses (bottom) in rewarded trials for increasing RT values in expert sessions (accuracy > 0.7) of one-sided mice split by trials with the associated or non-associated stimulus. Error bars indicate +/-s.e.m. (top, middle) and 95% confidence interval (bottom) of average across recordings. *p*-value calculated from two-sided t-test. **g**, Average DA stimulus-aligned signals (top), time warped signals (middle) and bar plots with average DA stimulus responses (bottom) across all mice for increasing psychometric slope and a fixed short RT range (0.5-2s) split by correct and error trials. **h**, Average DA stimulus-aligned signals (top), time warped signals (middle) and bar plots with average DA stimulus responses (bottom) across all mice for increasing RT values (same bins as in panel f) and a fixed large psychometric slope range (0.325-1) split by correct and error trials. * and ** indicate *p* < 0.05, *p* < 0.005 in all panels.

**Extended Data Fig.6.**
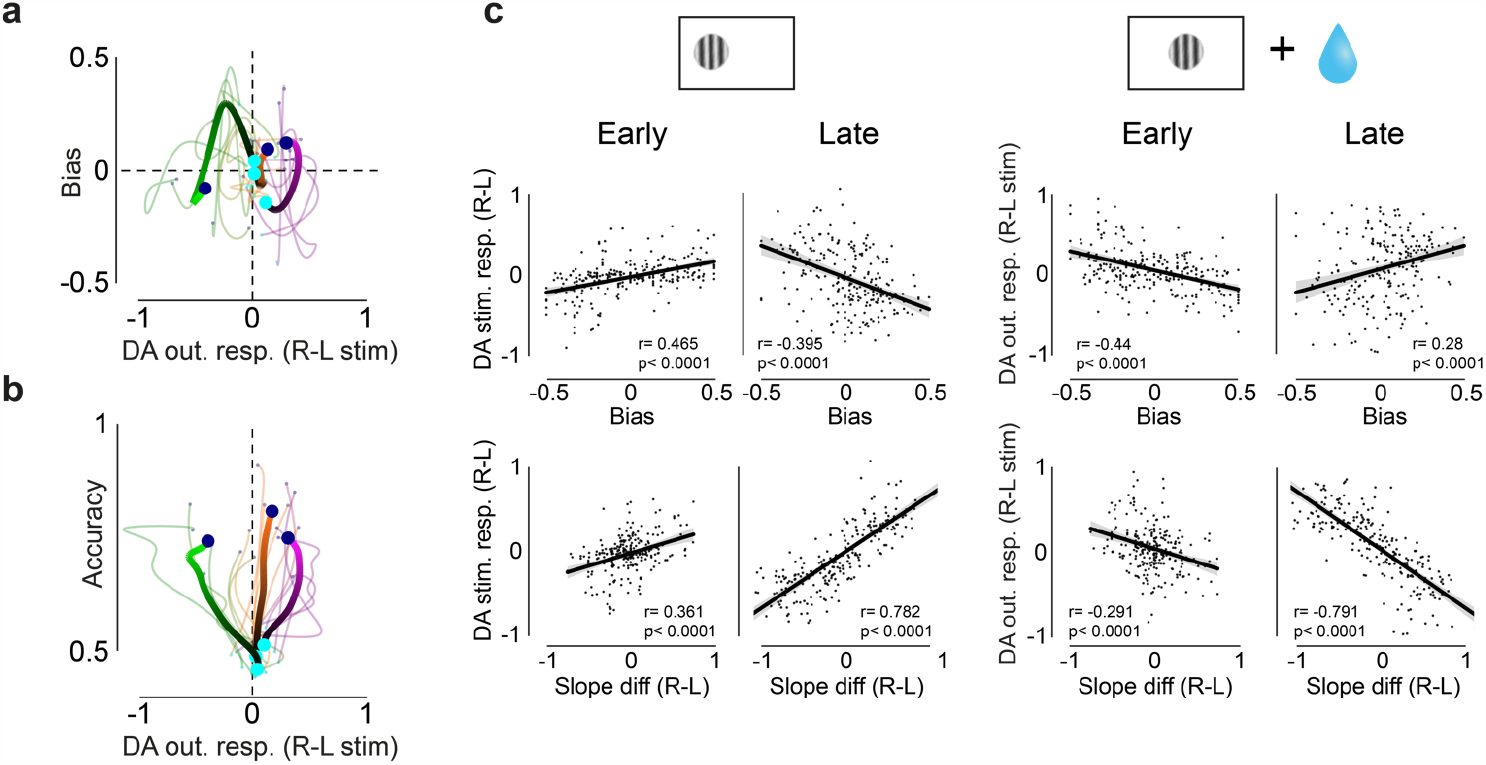
Further quantification of DLS DA responses to task events during learning. **a**, Difference in DA responses to rewarded outcomes after right and left stimulus (R-L) vs. bias, c.f. Fig.2i. **b**, Difference in DA responses to rewarded outcomes after right and left stimulus (R-L) vs. accuracy, c.f. Fig.2j. **c**, Left, regressions of early (accuracy not significantly greater than 0.5) and late (accuracy not significantly smaller than 0.7) bias and slope difference vs. difference in DA responses to right and left stimuli (R-L). Right, regressions of early and late bias and slope difference vs. difference in DA rewarded outcome responses after right and left stimuli (R-L). Each point represents a day. *p*-value calculated from the exact distribution of r.

**Extended Data Fig.7.**
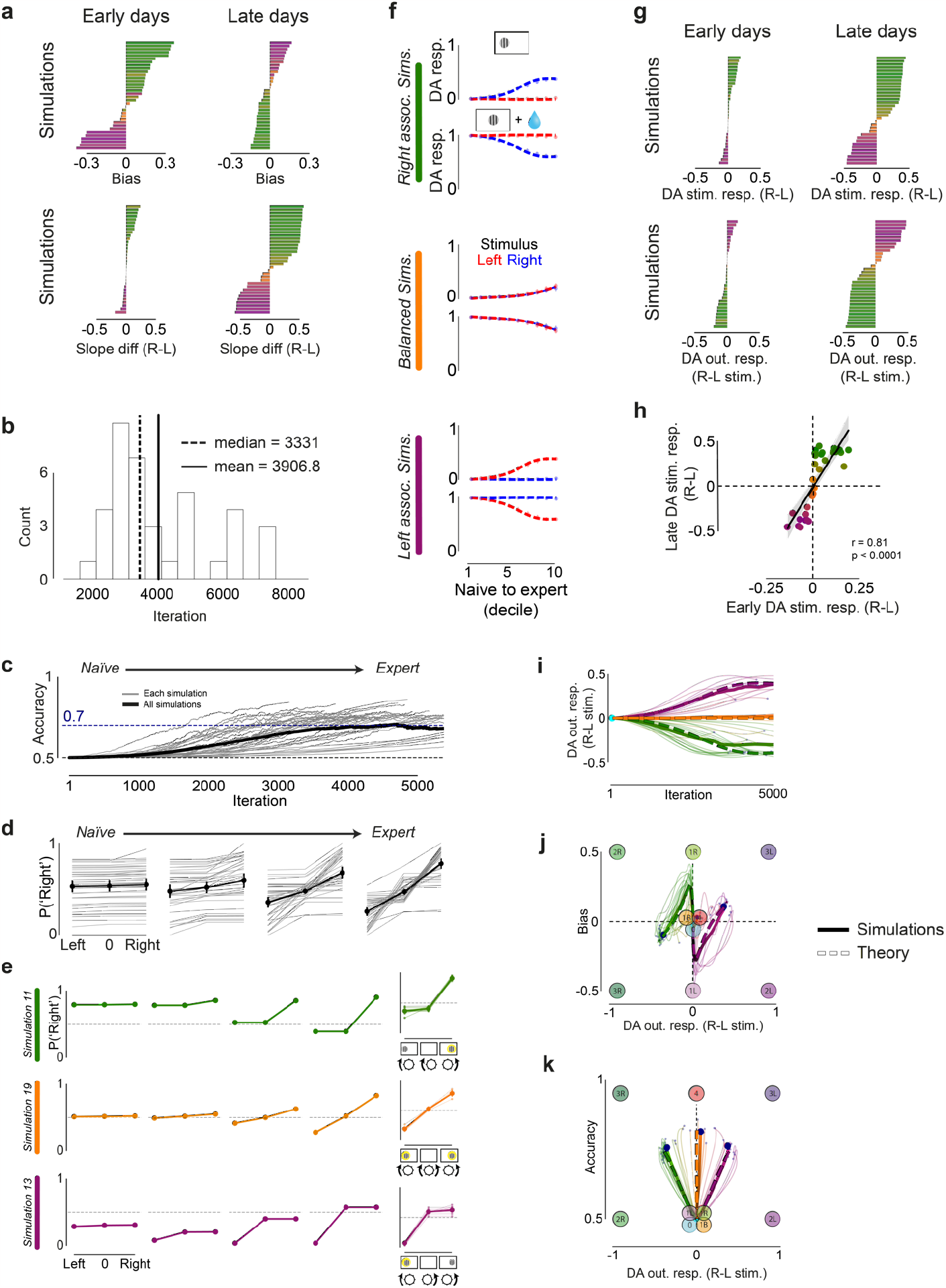
The tutor-executor network accounts for mouse behavioural trajectories and dopamine signals throughout learning. **a**, c.f. Extended Data Fig.2g, The model’s average bias and slope difference early in training (average across iterations 1000-2000) and late in training (average across final 1000 iterations) for each simulation. Colours come from average psychometric slope asymmetry of each simulation (see Extended Data Fig.2c). **b**, c.f. Extended Data Fig.1c, Histogram of the number of iterations that simulations required to reach 70% accuracy. **c**, c.f. Fig.1b, Accuracy over days per simulation (grey) and averaged across all simulations (black). Dashed lines indicate chance level (black) and 70% accuracy level (blue). Number of iterations limited to the average simulation length for better visualisation. **d**, c.f. Fig.1c, Psychometric curve over quartiles per simulation (grey) and averaged across all simulations (black). Error bars indicate 95% confidence interval across simulations. **e**, c.f. Fig.1d, Left, psychometric curves from 3 example simulations over quartiles throughout learning. Right, per simulation (thin) and average expert psychometric curves clustered by trajectory type (thick): right-associating (green), balanced (orange) and left-associating (purple). Cluster labels for each simulation obtained from Fig.3f, colours obtained from average psychometric slope asymmetry (see Extended Data Fig.2c). Error bars indicate 95% confidence interval of the average across simulations in each cluster. **f**, c.f. Fig.2d, Average model-derived stimulus and outcome DA responses over deciles in correct trials with stimulus on the left (red) and right (blue) for the three clusters from Fig.3f. Error bars indicate 95% confidence interval of average across simulations. Data points fit with a 3rd degree spline to visualise trends (scipy.interpolate.UnivariateSpline). **g**, c.f. Extended Data Fig.4h, the average difference in model-derived DA responses to right and left stimuli (R-L) and difference in model-derived DA rewarded outcome responses after R-L stimuli early in training (average across iterations 1000-2000) and late in training (average across final 1000 iterations) for each simulation. Colours come from average psychometric slope asymmetry (see Extended Data Fig.2c). **h**, c.f. Extended Data Fig.4g, Regression of early difference in model-derived DA responses to R-L stimuli (average across iterations 1000-2000) against late difference in model-derived DA responses (average across final 1000 iterations). Each point represents a simulation. *p*-value is calculated from the exact distribution of r. **i**, c.f. Extended Data Fig.4i, Difference in model-derived DA responses to rewarded outcomes after R-L stimuli over iterations per simulation (thin) and for the 3 clusters from Fig.3f (thick). Number of iterations limited to 5000 for better visualisation. **j**, c.f. Extended Data Fig.6a, Difference in model-derived DA responses to rewarded outcomes after R-L stimulus vs. bias. Stationary points here and in panel k are plotted using the average behaviour and DA predictions arising from their weight configurations. **k**, c.f. Extended Data Fig.6b, Difference in model-derived DA responses to rewarded outcomes after R-L stimulus vs. accuracy.

**Extended Data Fig.8.**
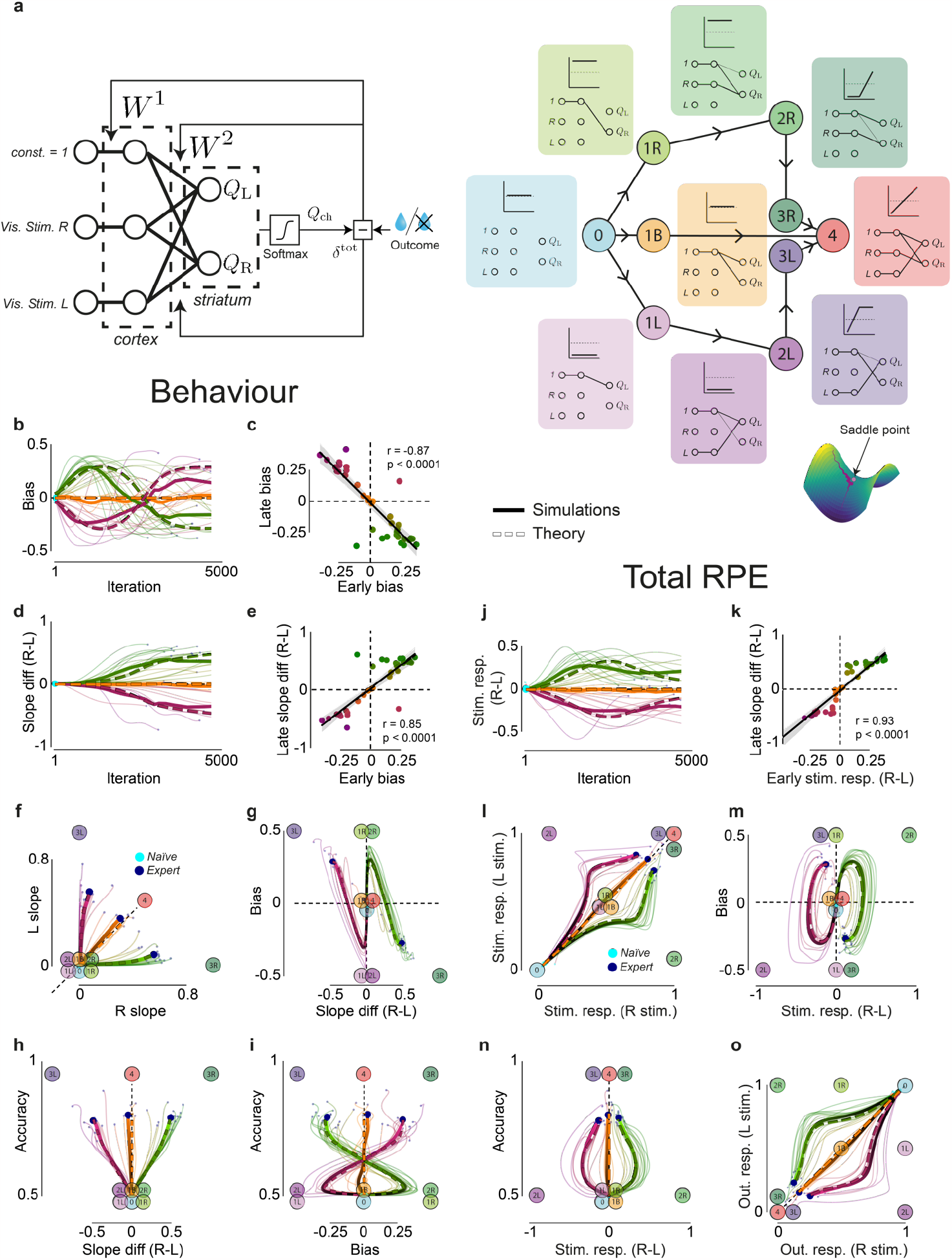
Single-loss gradient descent captures behavioural trajectories with a learning signal different from DLS DA. **a**, Left, schematic of the deep linear ‘single-loss’ RL network architecture and learning rule (Methods). Right, schematic of the stationary point structure with behavioural predictions as well as corresponding network weight configurations. The connecting lines with arrows represent the steepest heteroclinic orbits into/out of each stationary point (Methods). All the stationary points are saddle points except for 4, which is the global minimum. **b**, c.f. Fig.1f, Bias over iterations per simulation (thin), for the 3 clusters from panel f (thick), and for the average dynamics (thick dashed). Here, and in panels d and j, the number of iterations is limited to 4667 for better visualisation. Thick dashed lines in all panels indicate analytical trajectories derived from the average dynamics (Methods). **c**, c.f. Fig.1g, Regression of early bias (average across iterations 1000-2000) against late bias (average across final 1000 iterations). Each point represents a simulation. *P*-value calculated from the exact distribution of r. **d**, c.f. Fig.1h, Difference between right and left psychometric slopes over iterations per simulation (thin) and for the 3 clusters from panel f (thick). **e**, c.f. Fig.1i, Regression of early bias (average across iterations 1000-2000) against late slope difference (average across final 1000 iterations). Each point represents a simulation. *P*-value calculated from the exact distribution of r. **f**, c.f. Fig.1j, Left vs. right slope over iterations per simulation (thin) and for 3 clusters (thick). Clusters and colours obtained using the same procedure as for the behavioural data in Fig.1j. The clusters from this analysis are used in all other panels. Stationary points here and in panels g-I are plotted using the average behaviour arising from their weight configurations. **g-i**, c.f. Fig.1k-m, in order, difference in right and left (R-L) slope vs. bias, R-L slope vs. accuracy and bias vs. accuracy over iterations per simulation (thin) and for the 3 clusters from panel f (thick). **j**, c.f. Fig.2f, Difference in total RPE signals after right and left stimuli (R-L) over iterations per simulation (thin) and for the 3 clusters from Fig.3f (thick). **k**, c.f. Fig 2g, Regression of early stimulus-evoked total RPE (average across iterations 1000-2000) against late slope difference (average across final 1000 iterations). Each point represents a simulation. *p*-value calculated from the exact distribution of r. **l-o**, c.f. Fig.2h-k, in order, right vs. left stim.-evoked total RPEs; difference in total RPE evoked by right and left stimuli (R-L) vs. bias; difference in total RPE evoked by right and left stimuli (R-L) vs. accuracy; and outcome-evoked total RPE signals after right stimulus vs. after left stimulus per simulation (thin) and for the 3 clusters from Fig.3f. Stationary points are plotted using the average total RPEs arising from their weight configurations.

**Extended Data Fig.9.**
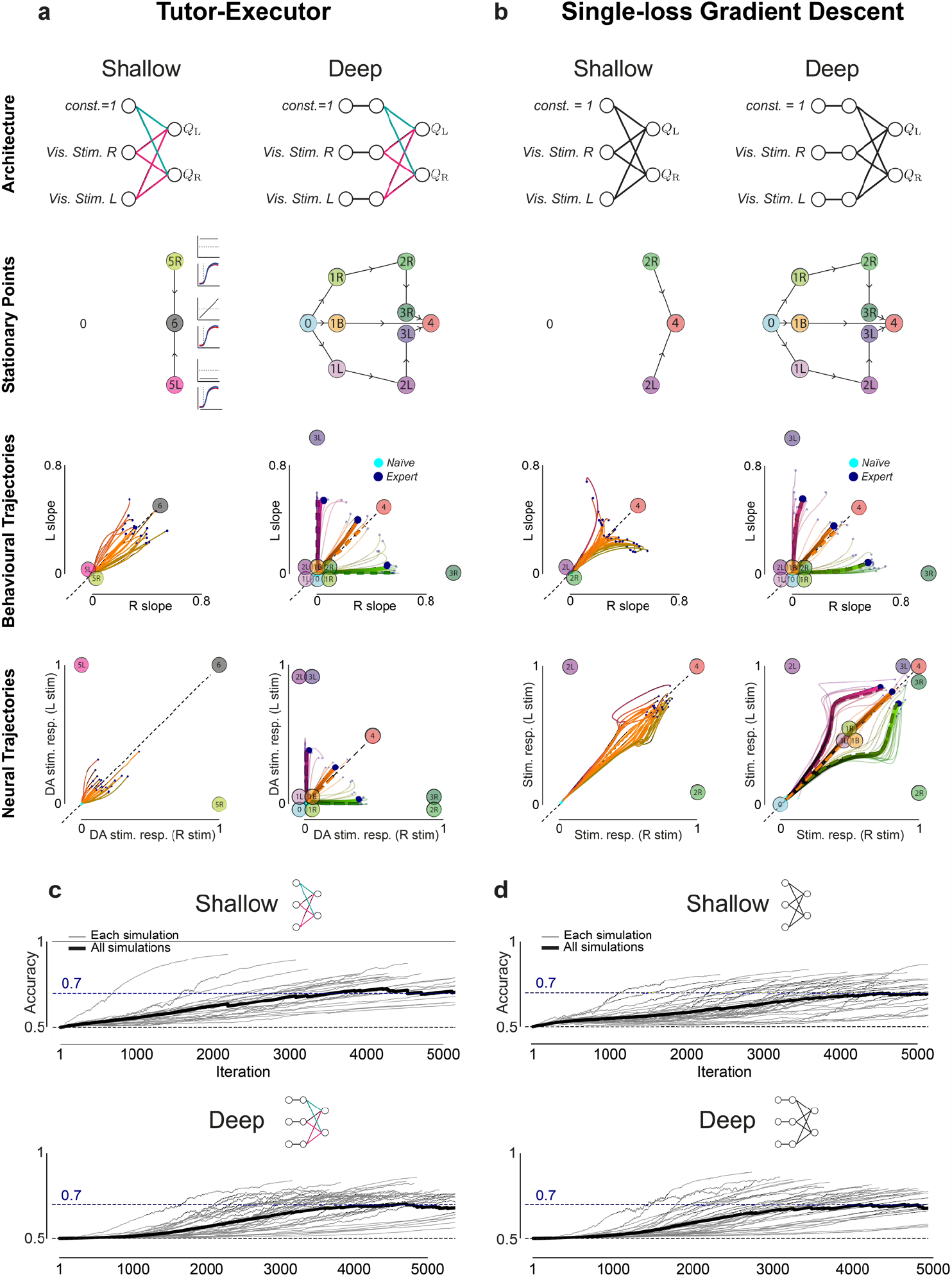
Depth is required for the model to capture the mice’s learning trajectories. **a**, Comparison between the architecture, stationary points, and simulated behavioural and neural trajectories (row for each) of a shallow (left column) and deep (right column) version of the tutor-executor model. Note that *0* is not a stationary point of the shallow network, and is placed in the left panel of the second row for reference. The shallow network has three stationary points which have different network configurations to those of the deep model. They are hence labelled with the increasing numbers *5L, 5R* and *6*, accompanied by schematics of their associated behaviour and neural predictions. These correspond to states where the shallow network is only making left choices (*5L*), right choices (*5R*) and the global optimum (*6*). **b**, Comparison between the architecture, stationary points, behavioural and neural trajectories (row for each) of a shallow (left column) and deep (right column) version of the single-loss gradient descent model. **c**, Comparison of the accuracy over iterations for simulations from a shallow (top) and deep (bottom) version of the tutor-executor model. Thin grey lines show the accuracy curves for each simulation, and the thick black line indicates the average across simulations. **d**, Comparison of the accuracy over iterations for simulations from a shallow (top) and deep (bottom) version of the single-loss gradient descent model. The learning curve of both deep models better captures mice data (c.f. Fig.1b) than the shallow models.

**Extended Data Fig.10.**
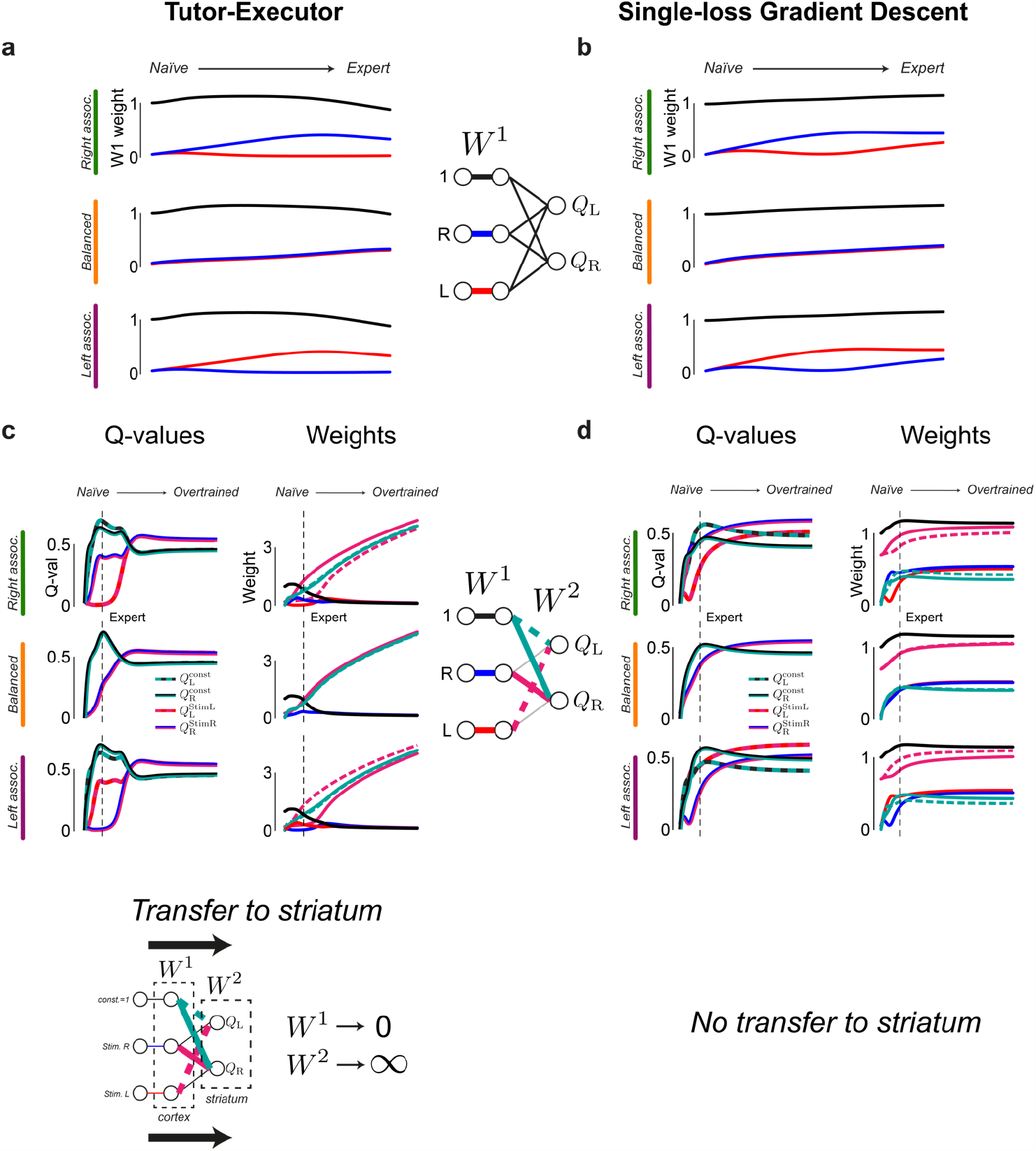
Tutor-executor learning rule causes transfer to striatum. **a**, *W*^*1*^ weights from naïve to expert in examples of right- (top), balanced (middle) and left-associating (bottom) analytical average dynamics of the tutor-executor network (same as dashed trajectories in Fig.3). Right- and left-associating trajectories were obtained by initialising the average dynamics with a small degree of left and right bias, whereas the balanced trajectory comes from a network initialised without bias (Methods). **b**, *W*^*1*^ weights from naïve to expert in examples of right- (top), balanced (middle) and left-associating (bottom) analytical average dynamics of the single-loss gradient descent network (same as dashed trajectories in Extended Data Fig.8). **c**, Left, stimulus and constant pathway Q-values (derived from the product of elements in *W*^*1*^ and *W*^*2*^, see Methods) from naïve to overtrained in examples of right- (top), balanced (middle) and left-associating (bottom) analytical average dynamics of the tutor-executor network. Overtrained: trained 8 times longer than the training used for naïve to expert. We do not plot the Q-values of the ‘incorrect’ associations (i.e., left stimulus with right choice and right stimulus with left choice) as these remain around 0. Right, *W*^*1*^ and *W*^*2*^ weights from naïve to overtrained. Again, we do not plot the weights that connect the inputs with the wrong choices as these remain around 0. **d**, similar to c but for the single-loss gradient descent model.

